# Stress, Heartbeats, and Biases: Cardiac phase and acute stress modulate affective and spatial response bias

**DOI:** 10.64898/2026.10.18.759627

**Authors:** Leon von Haugwitz, Romina Sadr, Edmund Wascher, Mauro F. Larrá

**Affiliations:** IfADo - Leibniz Research Centre for Working Environment and Human Factors, Ardeystraße 67, 44139 Dortmund, Germany

**Keywords:** Affective Priming, Interoception, Cardiac Cycle, EEG, Stimulus-Response Compatibility

## Abstract

Threatening situations often require rapid decisions which are particularly susceptible to response bias. The perception of threat also elicits an acute stress reaction, assumed to promote automatic action tendencies via several mechanisms. Cardioafferent signals during systole, for example, increase racially biased misidentifications of tools as weapons. Acute stress, linked to threat-biased decisions may both bias such decisions directly and alter cardiac effects on them. Here, we investigated the effects of cardiac phase and acute stress in a combined Weapon Identification and Simon Task. Thirty-one participants classified laterally or vertically presented weapons and tools with lateralized responses, such that target location and prime emotion provided task-irrelevant spatial and affective information. Target onsets were timed to systole or diastole, and in separate sessions participants were exposed to repeated feet cold pressor tests (CPT) and a warm-water control condition. The CPT reliably increased subjective, cardiovascular, and endocrine stress measures. Both systolic target onsets and CPT exposure strengthened the influence of emotional primes on weapon and tool classification, without evidence for general threat prioritization. For spatial compatibility, acute stress enlarged the Simon effect in sensitivity among fast responses, consistent with stronger initial automatic activation, and cardiac phase modulated the advantage of spatial correspondence over locations without dimensional overlap. Midfrontal theta showed prime-dependent target modulations with distinct patterns for cardiac phase and stress. Cardiac phase and stress did not interact, consistent with independent contributions rather than amplification or attenuation. Together, the findings indicate that acute stress and phasic cardioafferent signals both increase the influence of task-irrelevant affective and spatial information on action through partly distinct processes.

## 1. Introduction

Rapid decisions are easily biased by action tendencies elicited by features such as the location of a stimulus or the social or emotional meaning of a face. These automatically activate corresponding response tendencies that may facilitate or interfere with the intended action (De Houwer & Eelen, 1998; Hommel, 2011; Payne, 2001). How strongly such tendencies bias behavior also depends on the state of the body, whose signals shape brain function on timescales ranging from phasic fluctuations within each heartbeat (Azzalini et al., 2019; Critchley & Harrison, 2013; Engelen et al., 2023) to sustained states such as acute stress (Hermans et al., 2014). Phasic cardioafferent signals arise from arterial baroreceptors, whose firing increases during ventricular systole and decreases during diastole (Rau & Elbert, 2001). These signals reach the nucleus of the solitary tract via the glossopharyngeal and vagus nerves and are relayed from there to limbic and cortical regions including the amygdala, insula, and anterior cingulate cortex (Critchley & Harrison, 2013; Garfinkel & Critchley, 2016). Systolic stimulus onsets have typically been found to dampen sensory processing (Al et al., 2020; Edwards et al., 2007; Leupin & Britz, 2024; Salomon et al., 2018; Salomon et al., 2016), with threat-related stimuli as the notable exception (Garfinkel & Critchley, 2016). Fearful faces presented during systole are detected more easily, rated as more intense, and evoke larger amygdala responses than those presented during diastole (Garfinkel et al., 2014), and cardiac phase modulates attentional engagement with fearful faces and threat-related decisions (Azevedo et al., 2018; Mizuhara & Nittono, 2023; Yang et al., 2023). In their influential study, Azevedo et al. (2017) showed that cardiac phase can also modulate the expression of racial stereotypes. In the Weapon Identification Task (WIT), in which participants classify objects as weapons or tools following task-irrelevant face primes (Payne, 2001), Black relative to White face primes presented during systole increased the misidentification of tools as weapons, a finding since replicated (Sherrill et al., 2023). Notably, Azevedo and colleagues pointed out the potential relevance of this work for split-second decisions in police confrontations, a real-life instance of acute stress (Anderson et al., 2002; Baldwin et al., 2019; Bertilsson et al., 2019), yet it remains open how acute stress itself shapes the influence of affective information on such decisions and whether it alters cardiac-phase effects on them.

Acute stress engages the sympathetic nervous system within seconds and the hypothalamic–pituitary–adrenal axis, whose glucocorticoid output peaks tens of minutes later (Ulrich-Lai & Herman, 2009). The released mediators, including noradrenaline, corticotropin-releasing hormone, and cortisol, act on the brain in distinct but overlapping time windows (Joels & Baram, 2009), producing a sustained state that outlasts the stressor itself. This state is thought to redistribute processing from executive-control systems toward networks prioritizing salient information and rapid responding (Arnsten, 2009; Hermans et al., 2014), and acute stress alters neural responses to salient and emotional stimuli (Everaerd et al., 2017; Henckens et al., 2016; van Marle et al., 2009, 2010). In police officers, larger cortisol responses to a laboratory stressor were associated with fewer errors when deciding to shoot armed Black relative to armed White targets, suggesting heightened vigilance for stereotype-congruent threat (Akinola & Mendes, 2012), and high cortisol reactivity has been linked to reduced control over stereotypic responding (Amodio, 2009). Officers performing under anxiety likewise showed a response bias toward shooting, which the authors attributed to decisions based on threat-related expectations rather than task-relevant visual information (Nieuwenhuys et al., 2012). However, the behavioral consequences of acute stress for threat processing are inconsistent across paradigms (Andreotti et al., 2015; Hunter & Shields, 2023; Jiang et al., 2017; Metz et al., 2024), and whether stress changes how strongly task-irrelevant affective context, such as the emotional expression of a face, biases subsequent threat-related classifications remains unclear.

Acute stress might also change the cardiac modulation of affective processing, although the direction of such an interaction is difficult to predict. On the one hand, stress mediators exert excitatory effects on the amygdala (Joels, 2018; Joels & Baram, 2009; Ulrich-Lai & Herman, 2009), the structure through which systolic enhancement of fear processing has been proposed to operate (Azevedo et al., 2017; Garfinkel et al., 2014), so that stress could amplify cardiac effects. On the other hand, psychological stress suppresses baroreflex sensitivity (Gianaros et al., 2012), and the relative inhibition of fear processing during diastole is weaker in more anxious individuals (Garfinkel et al., 2014), so that stress could also attenuate them. Studies combining cardiac-cycle paradigms with a Cold Pressor Test (CPT) found no interaction in affectively neutral tasks (Mussini et al., 2024; von Haugwitz et al., 2024, 2026), but whether concurrent stress alters cardiac effects on affective processing, where both influences converge, has, to our knowledge, not been tested.

Prime effects in the WIT are analogous to affective compatibility effects, in which responses are facilitated when the valence of a task-irrelevant prime corresponds to that of the target or response (De Houwer & Eelen, 1998; Hermans et al., 1994; Herring et al., 2013). Such affective overlap is, however, not the only source of automatic response activation, and previous work from our group indicates that cardiac phase specifically enhances spatial activation in a Simon task (Larra et al., 2020; von Haugwitz et al., 2024). In the Simon task, the task-irrelevant stimulus location automatically activates the spatially corresponding response, producing better performance when stimulus and response locations match than when they conflict (Hommel, 2011; Simon, 1969). So called delta functions, that trace this compatibility effect across the entire RT distribution, show that this is particularly pronounced for fast responses and typically decreases toward the end of the response time distribution, consistent with a transient influence of stimulus-driven response activation (De Jong et al., 1994; Miller & Schwarz, 2021; Ridderinkhof, 2002). Presenting targets during systole selectively facilitated sensorimotor-compatible responses to lateralized auditory threat stimuli while impairing incompatible responses (Larra et al., 2020), a finding that was replicated for the standard visual Simon-task in a subsequent study (von Haugwitz et al., 2024). In a visual spatial-conflict task, cardiac-phase effects were associated with response-related rather than early sensory lateralization (von Haugwitz et al., 2026), consistent with an influence on response selection (Adelhofer et al., 2020). By contrast, although theoretical accounts predict increased stimulus-driven responding under stress, behavioral studies of conflict processing, including our own, have reported no stress effects or even improved performance (Duehnen et al., 2024; Plessow et al., 2011; Shields et al., 2019; von Haugwitz et al., 2024). In these paradigms, however, the response alternatives carried no affective meaning. It therefore remains open whether cardiac phase and acute stress act specifically on affective overlap or on automatic response activation more generally. It is equally open whether stress effects on spatial response activation emerge once spatial and affective overlap converge on the same response set.

We addressed these questions by combining an adapted WIT with a Simon task. Replacing the racial primes of the original task with fearful and neutral faces allowed us to test whether cardiac effects extend beyond racial stereotypes, as Azevedo et al. (2017) anticipated. Participants classified weapons and tools following these primes using lateralized responses, such that task-irrelevant target location induced spatial compatibility, whereas prime emotion could differentially bias weapon and tool classifications. Spatial and affective overlap thus acted on a single, affectively differentiated response set. In contrast to previous WIT studies, which timed prime onsets to the cardiac cycle (Azevedo et al., 2017; Sherrill et al., 2023), we timed target onsets to ventricular systole or diastole. Acute stress was manipulated within subjects using a repeated CPT and a warm-water control condition (Larra et al., 2015; Lovallo, 1975). Vertically presented targets provided a spatial non-overlap reference for separating effects associated with horizontal stimulus-response correspondence (Kornblum et al., 1990; Smith & Ulrich, 2025). In addition to response times and signal-detection measures, we examined compatibility delta functions to identify changes in early automatic response activation. We further assessed midfrontal theta activity as a marker of conflict-related control processes, which increases when automatically activated response tendencies have to be overridden (Asanowicz et al., 2021; Cavanagh & Frank, 2014; M. X. Cohen, 2014; Muralidharan et al., 2023; Tollner et al., 2017), and response-locked lateralized central beta power as an index of motor response selection (Asanowicz et al., 2021; Doyle et al., 2005; van Wijk et al., 2009). Together, the combined design allowed us to determine how cardiac phase and acute stress each shape the influence of task-irrelevant affective and spatial information on response selection, and whether the two interact.

## 2. Method

### 2.1. Sample

A total of 35 healthy individuals completed the experiment. Three participants were excluded from further analysis due to performance below chance level and one due to recording issues. The final sample comprised 31 participants (female = 15, age = 25.61 ± 3.84 years), all of whom were right-handed according to the Edinburgh Handedness Inventory (Oldfield, 1971).

Recruitment proceeded through social media channels and a university participant pool, so that the sample was drawn predominantly from the student population. Participants chose between course credit and payment (13.50 € per hour) as compensation. Eligibility was established in a screening interview conducted ahead of the appointment to include participants aged 18 to 35 years with a body mass index between 19 and 25 who reported good health both on the day of testing and throughout the two preceding weeks. Participants were excluded if they had taken medication or drugs within the preceding two weeks, apart from occasional over-the-counter preparations such as aspirin, if they reported any acute or chronic medical condition, in particular any indication of cardiovascular conditions or arrythmias, Raynaud’s disease, arterial hypertension, or a history of syncope, if hypertension was present in their family, or if they had a history of cerebral or aortic aneurysm or injury. Increased sensitivity to cold and complaints affecting oral health likewise precluded participation. Participants were further asked to refrain from caffeine, alcohol, exercise, and food during the two hours before each experimental session. The study received ethical approval from the IfADo ethics committee and adhered to the Declaration of Helsinki. Participants were briefed on the protocol and on their right to discontinue at any point before providing written informed consent.

### 2.2. Procedure

#### 2.2.1. General Procedure

Participants were seated in a dimly lit room in a comfortable chair, where EEG and ECG electrodes, a blood pressure cuff, and thoracic and abdominal respiratory belts were attached. After setup, participants completed two rating tasks of the stimuli used in the main task. First, they rated the face stimuli regarding their perceived facial expression on a 5-point Likert scale (dimensions: fear, disgust, joy, neutral, sadness, surprise, contempt, anger; cf. Langner et al. (2010)), which constitute the superset of expression categories across the three source databases (see 2.2.2.1). Second, participants classified the later target stimuli as weapon or tool, receiving corrective feedback after errors. They also rated each target in terms of identifiability, familiarity, perceived emotional valence, and threat. Participants then completed a training block of the experimental paradigm to familiarize themselves with the task. This was followed by a 10-minute resting phase. Subsequently, the first cold pressor test (CPT) or warm water control condition took place, immediately followed by the first experimental block. This procedure (CPT-Block) was repeated three additional times.

All experiments were conducted between 1:00 PM and 4:00 PM to account for diurnal variations in individual cortisol levels. Participants were neither informed about the scheduled condition (CPT vs. control) nor that it would alternate between sessions.

#### 2.2.2. Experimental task and apparatus

Participants performed a Weapon Identification Task (Payne, 2001) with an embedded Simon component (De Jong et al., 1994; Simon & Small, 1969) in which they had to classify a target object as a weapon or tool via left vs right button push after they were primed by either a fear or neutral face (Fig. 1 A). The target was presented at one of four equidistant positions (left, right, above, below) around central fixation with targets (i.e. weapon or tool) mapped to either left or right response buttons creating either spatially compatible, incompatible or neutral spatial conditions (Fig. 1 B). Specifically, trials were compatible when stimulus and response location matched (e.g. left target – left response), incompatible when a stimulus were presented opposite to the response location (e.g. right target – left response), or vertical/neutral when the target location was not represented in the response location (e.g. above target – right response). The response mapping was randomly counterbalanced between participants. Responses had to be made with the left and right index finger.

**Fig. 1.**
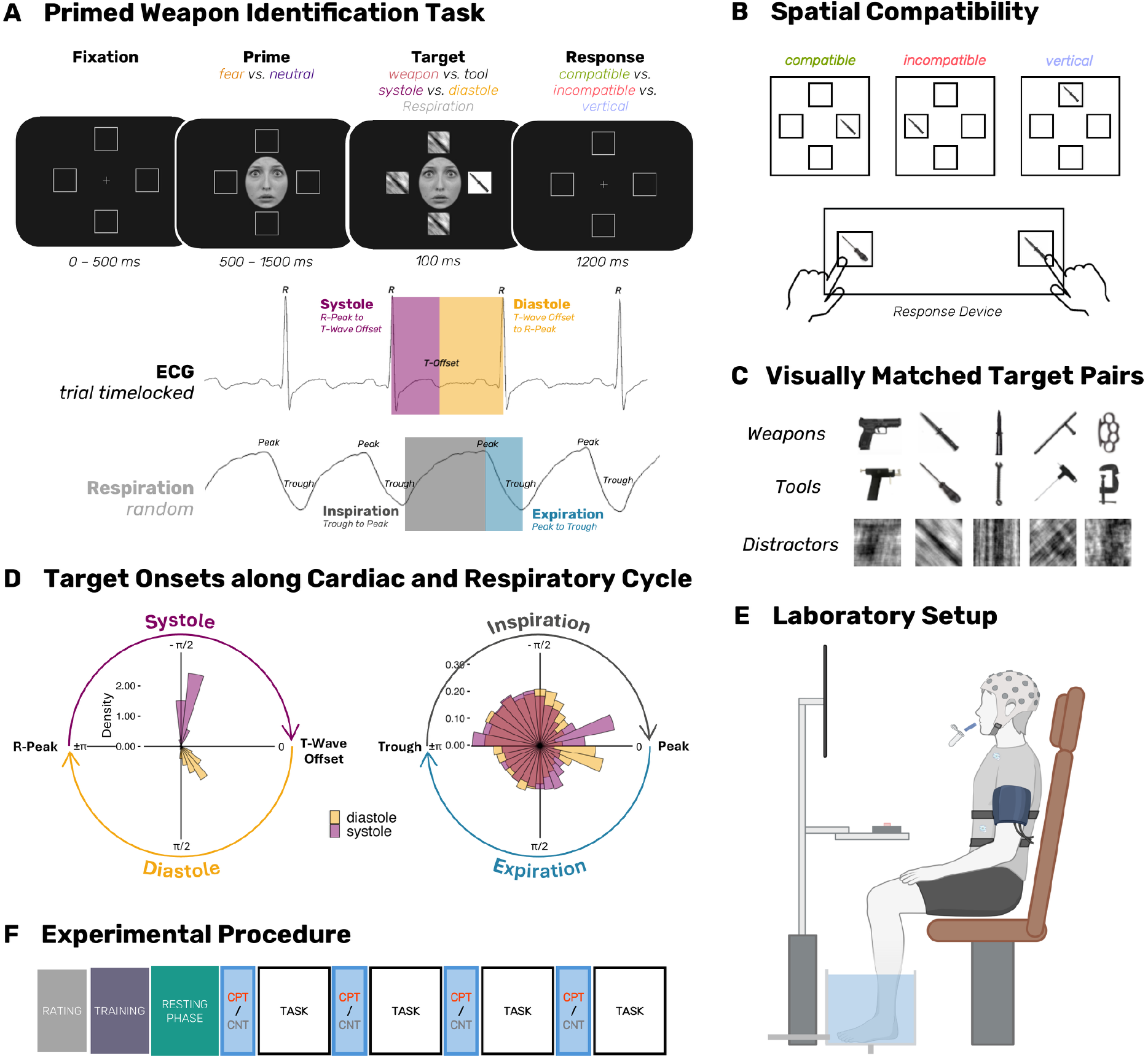
Schematic of joint Weapon Identification and Simon Task (Panel A) with possible spatial compatibilities (S-R, Panel B). Visually matched target pairs with fourier phase scrambled distractors based on pair specific merges (Panel C). Circular histograms of target onsets (Panel D) expressed as cardiac (left) and respiratory angle (right) as well as the laboratory setup (Panel E; created in BioRender. von Haugwitz, L. (2026) https://BioRender.com/ogyyxfn), and experimental procedure (Panel F).

During the experiment, participants were seated 70 cm in front of a 144 Hz monitor with a resolution of 1920 × 1080 pixels (Fig. 1 E). The response box was placed on a table in front of the participants at their midsagittal plain at a distance of approximately 30 cm and was of rectangular shape (13 cm × 37 cm) where the buttons were equally distanced from the center (10 cm). In each trial, a gray fixation cross (0.34° × 0.34°) was displayed at the center of the screen (viewing distance 80 cm). The fixation cross was surrounded by 4 gray squares (1.5° × 1.5°) presented equidistantly from the center on the left and right horizontal, and upper and lower vertical meridian at an eccentricity of approximately 1.39° representing the possible stimulus locations. Fixation cross and the squares were displayed continuously during the experimental blocks. After a jittered duration of 500-1000 ms, a centrally presented face stimulus was displayed for a jittered duration (500-1500 ms, approx. 61 px × 25 px, 1.34 ° × 0.55 °). Subsequently, one target and three distractor stimuli were presented for 100 ms each at the centers of the gray squares. The target was a single weapon or tool image whereby the remaining three locations were occupied by phase-scrambled filler stimuli matched to the target pairs in luminance, contrast, and spatial-frequency content (see Section 2.2.3).

Participants completed a training block of 64 trials followed by four experimental blocks of 256 trials each. Within each block, trials were balanced across prime emotion (fearful, neutral), target category (weapon, tool), target location (left, right, top, bottom), and cardiac phase (systole, diastole). Horizontal target locations (left, right), which carried the stimulus–response compatibility manipulation, were sampled twice as often as vertical locations (top, bottom). Trial order was pseudo-randomized separately for each participant and block under two sequence constraints: the same prime emotion appeared no more than three consecutive times, and the same face image was never presented on two consecutive trials.

Target onsets were presented time-locked to the cardiac cycle, either during systole (approx. at 160 ms) or diastole (approx. at 460 ms) after the R-wave peak (described in detail in section 2.2.4). Stimulus timing was managed using E-Prime 3.0 (PST Software, Inc.), with synchronization achieved through TTL pulses sent from an AccuSync 72 ECG monitor (AccuSync Medical Research Corporation) via serial interface.

##### 2.2.2.1. Face Primes and Target Stimuli

Face primes were taken from the Radboud Faces (Langner et al., 2010), NimStim (Tottenham et al., 2009), and the Karolinska Directed Emotional Faces database (Lundqvist et al., 1998), and comprised frontal-view, direct-gaze photographs of Caucasian female and male models displaying fearful and neutral expressions. Each identity contributed one fearful and one neutral image, yielding 24 female and 24 male identities (i.e., 48 fearful and 48 neutral primes). All faces were converted to grayscale and equated in mean luminance and contrast, and uninformative noise (e.g. hair) were cut-out to homogenize the faces and reduce interstimulus perceptual variance. This cutout–grayscale format has been shown to yield the most pronounced early emotion-sensitive responses without interacting with emotional expression itself (Schindler et al., 2019).

Targets were photographs of different weapons and everyday tools. Each weapon was assigned to a tool to form a matched shape pair, and within every pair both images were linearly rescaled to the pair’s mean luminance and root-mean-square contrast, so that a weapon and its paired tool were equated on these low-level properties. This procedure yielded 6 weapon and 6 tool targets. Filler stimuli that were presented alongside targets were generated by first merging the matched weapon and tool pair to one picture and subsequently fourier phase scrambling this merged image (Fig. 1 C; Oppenheim & Lim, 1981). This preserves the global luminance, contrast, and spatial-frequency content of the targets while abolishing recognizable form.

##### 2.2.2.2. Trial Onset along Cardiac Cycle

Target onsets were presented time-locked to the cardiac cycle, either during systole (M = 163 ms after R-Peak; phase angle = −81.6 °, SD = 7.4) or diastole (M = 464 ms, phase angle = 62.6 ± 17.1°).

Prior to analysis, we confirmed correct target timing via multiple checks. First, trials where R-Peak timing was based on implausible inter-beat intervals (IBIs) were excluded. Specifically, for each trial, we calculated the mean IBI of either the preceding 5 trials (for trials 6 onward) or the following 5 trials (for the first 5 trials), considering only IBIs within a physiologically plausible range of 400 – 1700 ms. Trials were excluded if their IBI deviated by more than 30% from this rolling average or fell outside the physiological range (M = 0.48 %, SD = 0.62; CPT: M = 0.48%; Control; M = 0.48%; t(30) = −0.08, p = .939). Second, we assessed the plausibility of systole duration (R-peak to T-wave offset) for each trial. Systole durations were flagged as implausible if they fell outside the physiological range of 100 - 400 ms or exceeded 3 median absolute deviations (MAD) from the subject-session median. For trials with implausible systole durations, we imputed the systole duration using a hierarchical imputation approach. Specifically, we first used the subject-condition mean of plausible systole durations, then the systole duration condition mean across all subjects, or finally the global mean of all plausible systole durations. Original systole durations were retained for 92.4% of CPT and 92.1% of control condition trials, with the remainder imputed predominantly from the subject’s session (CPT 8.4%, Control 8.8%).

Based on these cleaned systole durations, we computed cardiac angle following the procedure described by Sherman et al. (2022). Subsequently, we verified the accuracy of cardiac phase assignment by comparing intended phase (systole vs. diastole) with the adjusted phase calculated based on the cleaned timings. Trials whose intended cardiac phase onset did not match this timing-based classification were removed (M = 0.27 %, SD = 0.52 %; CPT: M = 0.26%; Control: M = 0.28%; t(30) = −0.11, p = .916).

#### 2.2.3. Stress Intervention: Automatic Cold Pressor Test

The bilateral cold pressor test (Larra et al., 2023; Larra et al., 2015) followed standardized protocols with automatically controlled timing, duration, and temperature via MATLAB. The water was cooled or heated to 3-4 °C (CPT) or 36-37°C (body temperature control), respectively, in shielded tanks with automatic continuous temperature control positioned in a separate room. Immersion occurred in rectangular basins (35 × 20 × 19 cm) using a rapid-fill/drain system (<8 s), with continuous circulation to prevent thermal stratification at the skin interface. Real-time monitoring ensured protocol fidelity via temperature/level sensors, basin-facing cameras to check submersion compliance, and automated temperature control (±0.5°C). Both conditions comprised 2-minute immersions during which participants were not informed about the remaining duration of exposure.

### 2.3. Data collection and analysis

#### 2.3.1. Saliva Cortisol

Saliva samples were collected using Salivettes (Sarstedt, Nürnbrecht, Germany). Samples were gathered after the first resting phase as a baseline, after each experimental block, as well as 30 min after the end of the experiment. Ahead of the experiment, participants were instructed to not touch the cotton swab, use their tongue to move it inside their mouth and to not chew on the cotton swab. Following the experimental sessions, the Salivettes were centrifuged, after which the samples were stored at −80°C until analysis. Free salivary cortisol was quantified after thawing by time-resolved immunoassay with fluorometric detection following the protocol described by (Dressendorfer et al., 1992).

#### 2.3.2. Subjective Ratings

Subjective stress and arousal were assessed after resting as baseline, immediately before and after every CPT or control exposure, and once more following the final experimental block. Both were indexed by visual analog scales (VAS) anchored at 0 (“not at all stressed/aroused”) and 100 (“extremely stressed/aroused”). The scales appeared on screen and participants positioned a cursor along the scale using the keyboard arrow keys.

#### 2.3.3. Mean Arterial Blood Pressure

Mean Arterial Blood pressure (MAP) was measured using a Dinamap Pro 300 V2 (Critikon; Tampa, Florida, USA) that was remote controlled via Eprime. Baseline blood pressure readings were taken at 3-minute intervals during the post-training resting phase. During the CPT and the warm water control procedure, blood pressure was measured 1 minute after water immersion.

#### 2.3.4. Heart Rate

Heart rate (HR) was continuously recorded using three electrodes in a modified Lead II chest configuration, connected to an AccuSync 72 ECG monitor (AccuSync Medical Research Corporation; Connecticut, Milford, USA). Offline artifact correction and R-peak detection were performed using *RDECO* (Moeyersons et al., 2019), with subsequent visual inspection to ensure accurate detection.

#### 2.3.5. Respiration

Respiration was measured with a thoracic and an abdominal belt and bandpass filtered from 0.05 to 3 Hz (second-order Butterworth, DC offset removed). To account for interindividual differences in abdominal and thoracic breathing, we combined the filtered thoracic and abdominal channels via their simple sum (Sola-Soler et al., 2023). The subsequent preprocessing was reimplemented in MATLAB following the rsp_process pipeline of NeuroKit2 (Makowski et al., 2021), which applies the extrema-detection approach of Khodadad et al. (2018). Peaks and troughs were identified by locating zero-crossings on the demeaned signal and finding local maxima and minima between crossings. Extrema whose vertical distance to a direct neighbor fell below 0.3 times the median vertical distance were discarded as outliers, and the sequence was sanitized to enforce alternating trough-to-peak-to-trough ordering. From the extrema we computed instantaneous respiration phase (inhalation, exhalation, undefined), a per-sample respiratory phase angle spanning −π to π across the breath cycle, instantaneous respiratory rate (breaths/min) from inter-peak intervals, and per-breath tidal amplitude. Note that rate was derived from inter-peak rather than inter-trough intervals, and the phase angle is expressed in radians analogue to cardiac phase angle. Beyond this pipeline, we rejected samples with low tidal amplitude (below 10% of the session median), implausible instantaneous rates (below 5 or above 40 bpm), and movement or spike events (thoracic or abdominal derivative exceeding 8 times the median sample-to-sample difference, extended by ±0.5 s).

#### 2.3.6. Manipulation Check Analysis

Salivary cortisol, subjective stress and arousal ratings, MAP, HR, and respiratory rate were analyzed with a common model. Each measure was entered into a linear mixed-effects (LMM) model fitted via the glmmTMB package (Brooks et al., 2017) in R (R Core Team, 2023) and the interaction of CONDITION (CPT vs. Control) with a categorical factor TIME, and a by-subject random intercept. Fixed effects were evaluated using Type III Wald chi-square tests via the car package (Fox & Weisberg, 2019), and the difference between CPT and Control was tested at each level of the time factor using estimated marginal means from the emmeans package (Lenth, 2025), corrected using the Benjamini-Hochberg procedure.

The levels of the TIME factor were measure specific. For heart rate and respiratory rate, they corresponded to normalized time bins. Each experimental phase (i.e. resting phase, CPT or Control procedure, experimental block) was divided into a fixed number of equal-width temporal bins. The number of bins per phase was determined a priori by dividing the median phase duration across participants and sessions by a target bin width of 120 s (the duration of the shortest phase, the CPT) and rounding to the nearest integer, yielding the same number of bins per phase for every participant. Within each participant and phase, bin width was then set to the phase duration divided by this fixed bin count, so that bins spanned the entire phase and represented equivalent normalized time points across participants despite minor between-participant differences in absolute duration. Mean heart rate was computed per bin from the artifact-corrected inter-beat intervals and mean respiratory rate from the cleaned instantaneous rate output. Cohen’s d was computed by dividing each contrast by the square root of the sum of the random-intercept and residual variances of the respective model (Westfall et al., 2014).

#### 2.3.7. EEG

EEG data were recorded using a 64-channel active electrode cap (actiCAP; Brain Products GmbH, Munich, Germany) positioned per the extended 10/20 system, sampled at 1 kHz. FCz served as the online reference, with AFz as ground. Electrode impedances were maintained below 15 kΩ using a BRAINAMP DC amplifier (Brain Products GmbH).

Preprocessing was performed in MATLAB R2024b with EEGLAB (v2025.0, Delorme & Makeig, 2004). Boundary events were removed, followed by bandpass filtering (0.1–40 Hz; 4th-order IIR Butterworth, DC offset corrected). Channels exhibiting flat signals (via clean_flatlines) or excessive noise were rejected using a multi-criterion approach: channels exceeding a z-score threshold of 3.29 (Rodrigues et al., 2021; Tabachnick & Fidell, 2007) for probability distribution, kurtosis, or spectral properties (1–125 Hz) were excluded. The union of channels flagged by these criteria was removed (M = 6.6 ± 1.0, range: 1-11). Missing channels were interpolated spherically (pop_interp), and data were re-referenced to the average.

Continuous data were epoched into 4-second segments time-locked to onset of the face primes (−1000 to +3000 ms). Trials with artifacts were iteratively rejected (pop_autorej; voltage threshold: ±1000 µV, probability threshold: 4 SD, max 5% rejection/iteration). Remaining data were downsampled to 250 Hz and bandpass-filtered (1-30 Hz; 4th-order IIR Butterworth). Independent component analysis (ICA; Infomax algorithm with sub-Gaussian source detection) was applied after adjusting component count to data rank via PCA. Components with <30% probability of neural origin (ICLabel; Pion-Tonachini et al., 2019) were rejected (mean: 24.45 ± 6.95, range: 10-42). Component labels and weights were mapped back to the original continuous dataset (1 kHz, interpolated, average-referenced).

Cardiac field artifacts (CFA) were removed in Python using the cfa_removal toolbox (Arnau et al., 2023; https://github.com/stefanarnau/cfa_removal). This neural network-based method predicts CFA from ECG using stimulus-free segments (−1000 to +1000 ms relative to R-peaks), then subtracts trial-specific CFA estimates from EEG.

Trials previously marked by autorej were rejected from this dataset, resulting in an average trial rejection rate of 11.28 ± 8.59 % (range = 1.81-69.9). For analysis of the WIT, the dataset was re-segmented into target locked (−800 ms to + 1200 ms) and response locked epochs (−1500 to +400 ms).

To reduce the spatial smearing introduced by volume conduction and to render the data independent of the recording reference (Kayser & Tenke, 2015), data was transformed into current source density (CSD) estimates using the spherical spline surface Laplacian (Perrin et al., 1989), as implemented in laplacian_perrinX (Mike X. Cohen, 2014).

##### 2.3.7.1. Event-Related-Spectral-Perturbations (ERSP) and Lateralized Power Spectra (LPS)

We examined two complementary spectral markers indexing different levels of processing. Midfrontal theta-band ERSP (4–8 Hz) served as an index of conflict monitoring and cognitive control demand (Asanowicz et al., 2021; Cavanagh & Frank, 2014; M. X. Cohen, 2014; Muralidharan et al., 2023; Tollner et al., 2017). Response-lateralized beta-band power (13–30 Hz) over central sites served as an index of motor-level response preparation (Asanowicz et al., 2021; Doyle et al., 2005; van Wijk et al., 2009).

Frequency components between 4 and 30 Hz were extracted in logarithmic steps via wavelet convolution, applied separately to each channel. The wavelet’s full width at half-maximum ranged from 750 ms at the lowest to 100 ms at the highest frequencies. Power was baseline corrected using the interval from −500 to −200 ms relative to target onset. Beta LPS was computed following Van der Lubbe and Utzerath (2013), analogously to the double-subtraction technique used for event related lateralization. Specifically, we subtracted ipsilateral from contralateral beta power at homologous central electrode pairs separately for left- and right-hand responses and scaled the average of both by the summed activation across hemispheres. Negative LPS thus indicates lower contralateral than ipsilateral power, that is, stronger contralateral suppression and stronger motor lateralization.

Statistical evaluation relied on cluster-based permutation tests using ft_freqstatistics (estimation method: Monte-Carlo; cluster correction: .05; cluster statistic: max t-sum; permutations: 10,000; two-sided) from the FieldTrip toolbox (Oostenveld et al., 2011). Midfrontal theta was averaged over a midfrontal region of interest (Fz, FCz, Cz) and over the theta band (4-8 Hz). For beta LPS, power was averaged across 13–30 Hz prior to clustering and tests were restricted to a central region of interest of nine homologous electrode pairs (FC1/FC2, FC3/FC4, FC5/FC6, C1/C2, C3/C4, C5/C6, CP1/CP2, CP3/CP4, CP5/CP6).

Because FieldTrip provides no factorial interaction test within this framework, interactions were evaluated by forming per-participant difference arrays for the contrast of interest at each fixed level of the remaining factors and submitting these to the same paired cluster test, so all interaction tests were pairwise and conditional rather than pooled. Since a non-significant cluster test cannot distinguish a true null from insufficient sensitivity (Cumming, 2014), we additionally report the mean per-subject difference (M) and its 95% confidence interval for every effect, over windows pre-specified from the expected time courses of theta engagement (100 to 600 ms post target onset, at the same ROI) and of beta LPS (−500 to 0 ms), i.e. fixed independently of the observed clusters (Kriegeskorte et al., 2009).

#### 2.3.8. Behavioral data

##### 2.3.8.1. Trial Rejection

Reaction times were screened on the log scale within each subject × session. Trials whose log-RT fell more than 3 MAD from the subject × session median were additionally flagged. This rejected 7.49 % (SD = 2.16) of trials per subject (CPT: M = 7.40, SD = 2.27%; Control: M = 7.58, SD = 3.20%; t(30) = −0.30, p = .766). Trials for which the respiratory phase could not be determined were removed (M = 6.63, SD = 6.18%; CPT: M = 7.11%; Control: M = 6.15%; t(30) = 1.25, p = .222). After all exclusion stages, on average 799.82 trials per subject and session were analyzed (SD = 74.69; CPT: M = 797.90 ± 74.58 trials; Control: M = 801.74 ± 75.98 trials; t(30) = −0.34, p = .738).

##### 2.3.8.2. Response Times and Signal Detection Measures

Response times and accuracy were analyzed using generalized linear mixed-effects models (GLMMs), again fitted in glmmTMB and evaluated with Type III Wald chi-square tests via car. The fixed effects structure of both models included all main effects and interactions among CONDITION (CPT vs. Control), PRIME (fear vs. neutral), TARGET (weapon vs. tool), COMPATIBILITY (compatible, incompatible, neutral), and CARDIAC PHASE at target onset (systole vs. diastole), with respiratory phase at target onset entered as a nuisance covariate via its sine and cosine components as a linear circular approximation. The random effects structure was determined through iterative model comparison using likelihood ratio tests (LRTs) following (Bates et al., 2015) and (Matuschek et al., 2017), starting from a maximal structure and simplifying where variance components approached zero or where the model failed to converge with a positive-definite Hessian. Both models retained a by-subject random slope for experimental condition, and a by-stimulus-item random intercept and slope for experimental condition were included for target stimuli (n = 10). Random effects for prime face identity (n = 96) were excluded in both models (RT: χ² < 0.001, df = 1, p = 1.00; SDT: χ² = 2.86, df = 1, p = .091, with BIC favoring the simpler model). Model assumptions and convergence were verified for both models using simulation-based residual diagnostics implemented in DHARMa (Hartig, 2016) based on optimizer exit codes, gradient norms at the optimum, and positive-definiteness of the Hessian matrix. Simple effects to decompose significant interactions were obtained using joint_tests() from the emmeans package (Lenth, 2025), evaluated separately at each level of the conditioning factor. Estimated marginal means (EMMs) were computed to examine significant effects, and pairwise comparisons were corrected for multiple testing using the Benjamini-Hochberg procedure.

Response times (RTs) from accurate trials were modelled with a gamma distribution with log link, with a by-subject random slope for target type additionally retained. Because within-group residuals were heterogeneous across levels of spatial compatibility, the final model included spatial compatibility as a predictor of the Gamma shape parameter.

Accuracy was analyzed within a signal detection theory (SDT) framework adapted for use with GLMMs (DeCarlo, 1998), which decomposes accuracy into sensitivity (d′) and response bias (c) without requiring prior aggregation across trials. Trial-level weapon responses (responded_weapon: 0/1) were modelled using a binomial distribution with probit link and stimulus type (weapon vs. tool) coded as is_signal (1/0), such that the is_signal coefficient and its interactions capture modulations of d′, while the intercept and remaining main effects capture shifts in response criterion. A by-subject random slope for is_signal was retained.

##### 2.3.8.3. Delta and Conditional Accuracy Functions

To examine how the compatibility effect varied across the RT distribution, we analyzed delta functions (De Jong et al., 1994), which plot the compatibility difference against RT and thereby trace the temporal course of the Simon effect. The negative-going slope commonly observed in the horizontal Simon task, for example, is assumed to represent initial response activation by the irrelevant target location followed by its later suppression (Miller & Schwarz, 2021; Ridderinkhof, 2002). Rather than computing RT quantiles within participant and condition and then taking pairwise differences between conditions at each quantile position, which discards trial-level information and would not allow us to control for respiratory phase, we estimated conditional quantiles of the RT distribution directly as a function of experimental predictors using trial-level quantile regression via the quantreg package (Koenker, 2025).

To address between-participant differences in RT distributional shape, which the two-step approach removes implicitly, each trial’s RT was replaced by its within-participant rank divided by (n + 1), where n is the participant’s total number of trials, mapping each participant’s RT distribution onto a uniform [0, 1] scale. Because this probability integral transform (PIT) is monotone but nonlinear, absolute delta function shape is not directly readable from the PIT scale. For that reason, the central shape claim of the present analysis, the negative-going slope of the Simon compatibility delta function, was verified independently on conventional Vincentized delta functions computed within participant on the millisecond scale (Ratcliff, 1979), displayed in the supplementary materials (Fig. S3). Importantly, comparisons between conditions are unaffected, because the transform is estimated once per participant over all trials, so at a given quantile every condition is weighted by the same density factor.

Quantile regression models were fitted at nine quantile positions (10%–90%) using the Frisch-Newton interior point method (method = “fn”) via rq(), with the fixed effects specified above. Compatibility was coded with treatment contrasts (compatible as reference), yielding three coefficients directly corresponding to the Simon compatibility delta (incompatible − compatible), a Simon facilitation delta (vertical − compatible, CV), and a Simon inhibition delta (vertical − incompatible, IV) (Smith & Ulrich, 2025). Standard errors were clustered on participant using the cluster-robust sandwich estimator for quantile regression (Parente & Santos Silva, 2016), with Wald statistics referred to t(G − 1) and F(q, G − 1) following Cameron and Miller (2015), where G = 31 is the number of participants. To test whether effects varied across the RT distribution we tested the null hypothesis that a given contrast is constant across all estimated quantile levels, using the eight differences between adjacent quantiles. Per-quantile contrasts were then examined with their cluster-robust standard errors, corrected across the nine quantiles within each contrast using the Benjamini-Hochberg procedure.

Conditional accuracy functions (CAFs) were estimated using the SDT GLMM described above but included within-subject PIT-transformed RT as a continuous predictor that interacted with all factors. RT was parameterized as a cubic polynomial, supported by a model comparison favoring the cubic polynomial over natural-spline parameterizations with four (ΔAIC = 55) and five (ΔAIC = 96) degrees of freedom, with visually checked goodness-of-fit (Fig. S1) and a five-fold subject-held-out cross-validation indicated good generalization (CV log-loss = 0.525 vs. in-sample log-loss = 0.517, a 1.6% increase). As in the quantile regression, two reference codings of COMPATIBILITY were used to recover all three pairwise delta functions. Via linearHypothesis() from car (Fox & Weisberg, 2019), we decomposed interactions involving COMPATIBILITY × RT into per-pairwise contributions (CI, CV, IV) via 3-df joint Wald χ² tests on the relevant subsets of polynomial × moderator coefficients, paralleling the slope-equality decomposition used for the quantile regression. Conditional d′ estimates were extracted at the same nine quantiles using EMMs, and p-values for the family of tau-specific compatibility contrasts were adjusted using the multivariate-t method, which provides family-wise error-rate control based on the joint covariance structure of the contrasts. We did not include criterion delta functions in our analysis, as we did not have specific hypothesis regarding their RT dependence and stress or cardiac phase modulations.

## 3. Results

### 3.1. Manipulation Check

The CPT induced the expected endocrine, cardiovascular, and subjective stress response (Figure 2). The LMMs (Table S1) revealed a significant CONDITION × TIME interaction for salivary cortisol (χ²(6) = 58.47, p < .001), MAP (χ²(8) = 229.82, p < .001), subjective stress (χ²(8) = 63.30, p < .001), arousal (χ²(8) = 102.94, p < .001), HR (χ²(41) = 209.01, p < .001), and respiration rate (χ²(41) = 65.60, p = .009). Conditions did not differ at baseline in any measure (all p ≥ .069). Cortisol was elevated in the CPT condition from the second post-block sample through the first post-experiment sample and had returned to control levels approximately 120 min after the start of the resting phase (Table S2). MAP was elevated during, and subjective stress and arousal after, each CPT (Tables S3–S5). MAP no longer differed between conditions after the experimental blocks, whereas subjective stress and arousal remained elevated after blocks 3 and 4. HR was elevated during each CPT but fell below control levels during large parts of all four experimental blocks (Table S6). Respiration rate was lower during the first three CPTs and did not differ between conditions during the experimental blocks (Table S7).

**Fig. 2.**
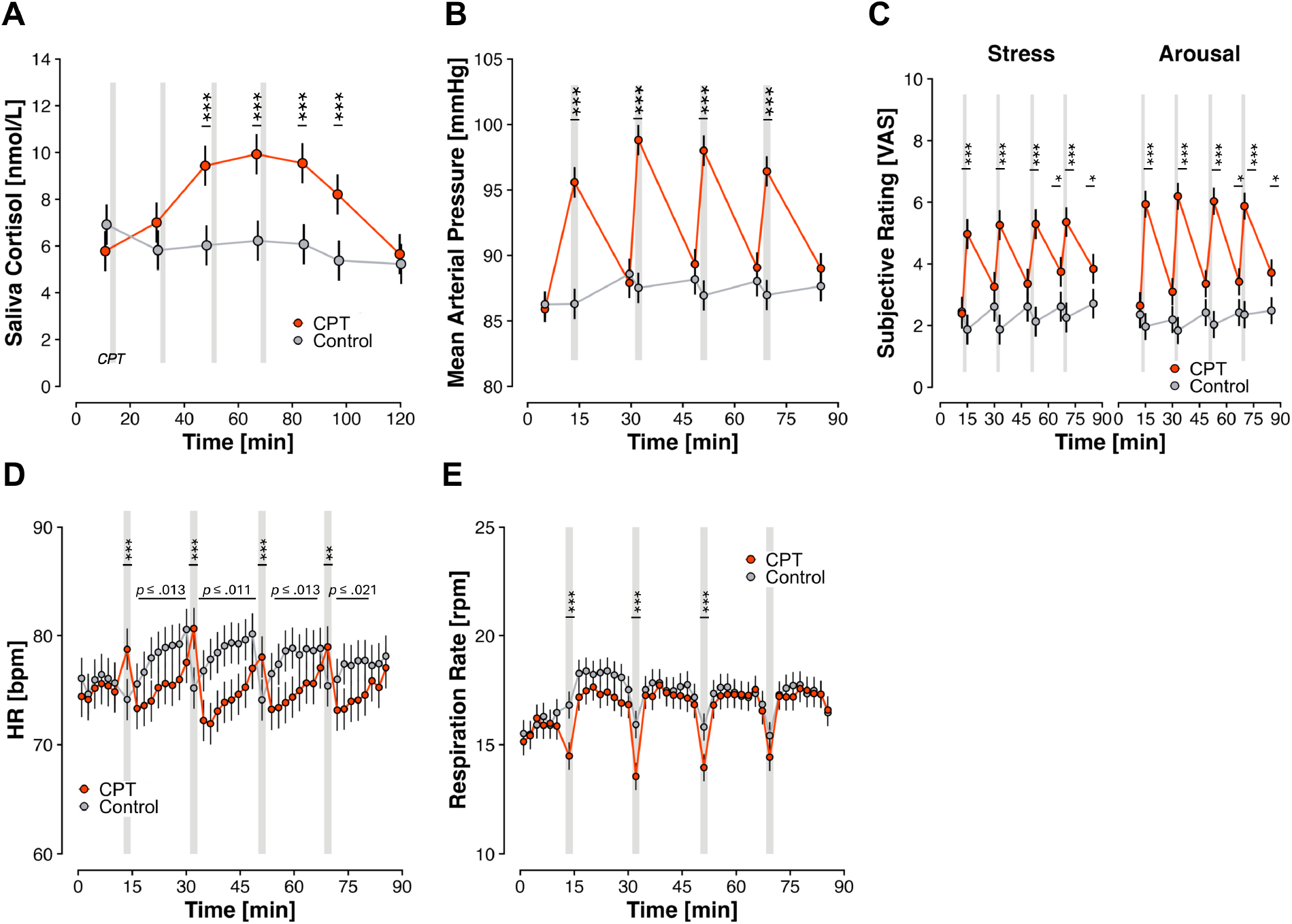
Saliva Cortisol (Panel A), Mean Arterial Blood Pressure (MAP, Panel B), Subjective Stress and Arousal (Panel C), Heart Rate (HR, Panel D), and Respiration Rate (Panel E) during the experiment for the CPT and control condition. Gray bars mark the timing of the CPTs or warm water control procedures. Error bars show standard errors. *\* p < .05, ** p < .01, *** p < .001*.

### 3.2. Behavioral measures

#### 3.2.1. Reaction Time

The GLMM on RT revealed significant main effects of COMPATIBILITY (χ²(2) = 96.93, p < .001) and RESPIRATION (sine; χ²(1) = 6.38, p = .012), as well as interaction effects of COMPATIBILITY × TARGET (χ²(2) = 14.16, p < .001), CONDITION × PRIME × TARGET (χ²(1) = 6.91, p = .009), and CARDIAC PHASE × PRIME × TARGET (χ²(1) = 4.72, p = .030).

As expected, the main effect of COMPATIBILITY confirmed that compatible trials showed significantly lower RTs versus incompatible (z = -4.92, p < .001, ΔRT = -25.04 ± 5.09) and vertical (z = -9.62, p < .001, ΔRT = -39.26 ± 4.14) trials, whereby vertical trials showed significantly higher RTs than incompatible trials (z = -2.53, p = .011, ΔRT = -14.23 ± 5.63).

This compatibility pattern was qualified by TARGET (COMPATIBILITY × TARGET). Simple effects analyses showed that an effect of TARGET emerged specifically on vertical trials (χ²(1) = 4.47, p = .035; comp: χ²(1) = 1.70, p = .192; incomp: χ²(1) = 0.83, p = .362), with faster responses to weapons than tools (z = 2.11, p = .035, ΔRT = 43.81 ± 20.70). Conversely, the structure of the compatibility effect itself differed between targets: for tool targets, all three pairwise contrasts were significant (comp vs. incomp: z = -3.67, p < .001, ΔRT = -21.22 ± 5.86; comp vs. vertical: z = -9.74, p < .001, ΔRT = -46.86 ± 4.74; incomp vs. vertical: z = -3.94, p < .001, ΔRT = -25.64 ± 6.39). By contrast, for weapon targets RTs on incompatible and vertical trials did not differ (z = -0.54, p = .590, ΔRT = -3.34 ± 6.20), while both were slower than compatible trials (comp vs. incomp: z = -5.11, p < .001, ΔRT = -28.70 ± 5.62; comp vs. vertical: z = -6.97, p < .001, ΔRT = -32.04 ± 4.48).

Follow-up analyses of the CONDITION × PRIME × TARGET interaction revealed multiple effects. First, we found a CPT specific interaction of PRIME × TARGET (χ²(1) = 11.94, p < .001) not present in the control condition (χ²(1) = 0.71, p = .398). This CPT specific interaction showed that effect of primes differed ordinally for weapons and tools (z = 3.17, p = .002, ΔRT = 10.9 ± 3.44). Specifically, fear versus neutral primes elicited faster responses for weapons (χ²(1) = 4.69, p = .030, z = -2.17, ΔRT = -5.21 ± 2.41) whereas for tools the opposite was the case, i.e. when neutrally primed participants showed faster responses to tools than when fear primed (χ²(1) = 5.37, p = .021; z = 2.32, ΔRT = 5.67 ± 2.45). Second, simple effect analyses showed a significant interaction of CONDITION × TARGET specifically for fear primes (χ²(1) = 4.65, p = .031; tools: χ²(1) = 0.03, p = .864). Post-hoc tests of this interaction indicated that the difference between tool and weapon targets was significantly higher in the CPT vs. control condition (z = 2.16, p = .031, ΔRT = 11.4 ± 5.77).

Simple effect analyses of the PHASE × PRIME × TARGET showed a significant PRIME × TARGET interaction specifically for systole onsets (χ²(1) = 8.06, p = .005; diastole: χ²(1) = 0.05, p = .817). Analogue to the CPT effect, the effect of primes differed ordinally for weapons and tools (z = 2.84, p = .005, ΔRT = 9.84 ± 3.47). Specifically, neutral versus fear primes evoked faster responses for tool targets (χ²(1) = 4.33, p = .038, z = 2.08, ΔRT = 5.14 ± 2.47). While not significant, descriptively fear versus neutral primes elicited faster responses for weapons (χ²(1) = 3.76, p = .053, z = -1.94, ΔRT = -4.71 ± 2.43).

Finally, the sine component of respiratory phase predicted RT (χ²(1) = 6.38, p = .012), whereas the cosine component did not (χ²(1) = 0.03, p = .856). Because the two components jointly encode a single circular predictor, we combined them in a 2-df Wald test of whether respiratory phase modulated RT overall which confirmed modulation across the respiratory cycle (χ²(2) = 6.40, p = .041), carried entirely by the sine component. As the sine term is negative throughout inhalation and positive throughout exhalation, RT tracked respiratory flow direction (inhale vs. exhale) rather than lung volume (full vs. empty), with RTs fastest near mid-inhalation and slowest near mid-exhalation with an average peak-to-trough difference of 4.50 ± 1.79 ms (cf. Supplementary Figure 2).

There were no further significant main or interaction effects on RT (TARGET: χ²(1) = 2.02, p = .155; PRIME × TARGET: χ²(1) = 3.40, p = .065; CARDIAC PHASE × RESPIRATION (sine): χ²(1) = 3.31, p = .069; CONDITION × TARGET: χ²(1) = 1.23, p = .268; CONDITION × PRIME × CARDIAC PHASE: χ²(1) = 2.51, p = .113; all other χ² < df).

#### 3.2.2. Signal Detection Measures: Sensitivity and Criterion

The SDT GLMM revealed effects on both sensitivity (d’) and criterion. As shown in Figure 4A, subjects discriminated weapon and tool stimuli well above chance (d′ = 2.02, z = 5.21, p < .001), with no significant overall response bias towards weapons or tools (c = −0.03, z = −0.18, p = .860, Figure 4F). Furthermore, both sensitivity and criterion were modulated by COMPATIBILITY (sensitivity: χ²(2) = 893.13, p < .001; criterion: χ²(2) = 198.45, p < .001), and an interaction of CONDITION × COMPATIBILITY (sensitivity: χ²(2) = 9.25, p = .010; criterion: (χ²(2) = 29.49, p < .001).

The COMPATIBILITY effect on sensitivity showed that sensitivity was significantly higher in compatible versus incompatible (z = 23.26, p < .001, Δd’ = 0.86 ± 0.04) and vertical trials (z = 28.71, p < .001, Δd’ = 1.06 ± 0.04), whereby incompatible trials showed significantly higher sensitivity than vertical trials (z = 5.74, p < .001, Δd’ = 0.20 ± 0.03).

As indicated by the COMPATIBILITY × CONDITION × d’ interaction this compatibility effect was modulated by CPT exposure. Post hoc contrasts revealed that the sensitivity difference between compatible and incompatible trials was significantly higher in the CPT compared to the control condition (z = 3.00, p = .008, Δd CPT vs. Δd Control = 0.22 ± 0.07) while CPT and control condition did not differ in Δ CV (z = 1.26, p = .207) or Δ IV (z = −1.87, p = .092). Simple effect analyses showed that sensitivity between CPT and control condition did not differ significantly in compatible (χ²(1) = 2.90, p = .088), incompatible (χ²(1) = 1.02, p = .312) or vertical trials (χ²(1) = 0.40, p = .529).

The COMPATIBILITY effect on criterion revealed that compatible trials had a significantly more conservative criterion than vertical trials (z = 3.05, p = .007; Δc = 0.15 ± 0.05), as did incompatible trials (z = 2.40, p = .024, Δc = 0.10 ± 0.04). Compatible and incompatible trials did not differ in criterion (z = 0.72, p = .471; Δc = 0.05 ± 0.07). However, none of the compatibility conditions individually showed a criterion that significantly deviated from zero, i.e. a significant response bias towards weapons or tools (compatible: z = 0.18, p = .858, c = 0.03 ± 0.19; incompatible: z = -0.08, p = .934, c = -0.02 ± 0.19; vertical: z = -0.62, p = .536, c = -0.11 ± 0.19).

The interaction of CONDITION × COMPATIBILITY showed that this compatibility pattern on criterion was again modulated by CPT exposure. Specifically, under control conditions vertical trials showed a significantly lower criterion versus incompatible trials, an effect that was not present in the CPT condition (Control: z = 3.61, p < .001, Δc = 0.16 ± 0.04; CPT Δc = 0.04 ± 0.04, z = 0.83, p = .405). In line, this ΔIV in criterion was significantly higher under control versus CPT condition (z = -3.67, p < .001, CPT vs. Control = -0.12 ± 0.03). While both under CPT and control conditions, the criterion between compatible and incompatible trials did not differ, this ΔCI was still significantly higher in the CPT compared to the control condition (z = 4.29, p < .001, CPT vs. Control: = 0.16 ± 0.04). Under both CPT and control condition, compatible trials showed a significantly more conservative criterion compared to vertical trials (CPT: z = 3.16, p = .002, Δc = 0.16 ± 0.05; Control: z = 2.56, p = .010, Δc = 0.13 ± 0.05). Direct comparisons between CPT and control condition did not reveal significant differences of criterion in compatible (χ²(1) = 1.51, p = .23), incompatible (χ²(1) = 1.80, p = .18), or vertical trials (χ²(1) = 0.54, p = .461). Finally, none of the six cell-level criteria differed significantly from zero (all z ≤ 0.68).

There were no further significant main or interaction effects on criterion or sensitivity (criterion: RESPIRATION (cos): χ²(1) = 2.77, p = .096; CONDITION × PRIME × COMPATIBILITY: χ²(2) = 4.80, p = .091; RESPIRATION (sin): χ²(1) = 2.03, p = .154; CONDITION × COMPATIBILITY × CARDIAC PHASE: χ²(2) = 2.38, p = .304; all other criterion χ² < df; sensitivity: CONDITION × PRIME × COMPATIBILITY × d’: χ²(2) = 2.58, p = .275; CONDITION × PRIME × d’: χ²(1) = 1.96, p = .162; CARDIAC PHASE × RESPIRATION (sin) × d’: χ²(1) = 1.69, p = .193; CARDIAC PHASE × d’: χ²(1) = 1.43, p = .231; PRIME × d’: χ²(1) = 1.21, p = .271; RESPIRATION (sin) × d’: χ²(1) = 1.16, p = .281; all other sensitivity χ² < df).

#### 3.2.3. Distributional Analysis of Compatibility Effects

##### 3.2.3.1. RT Deltas

As expected, the clustered quantile regression on PIT-transformed RT showed that all three COMPATIBILITY deltas varied in magnitude along RT (Δ CI: F(8, 30) = 7.63, p < .001; Δ CV: F(8, 30) = 23.94, p < .001; Δ IV: F(8, 30) = 3.82, p = .003). Furthermore, it indicated interaction effects of COMPATIBILITY × TARGET (Δ CV: F(8, 30) = 2.38, p = .041), COMPATIBILITY × CARDIAC PHASE (Δ IV: F(8, 30) = 2.88, p = .017), and COMPATIBILITY × CARDIAC PHASE × PRIME (Δ CV: F(8, 30) = 2.30, p = .047).

A linear-in-τ contrast computed from the clustered covariance of the quantile process showed that the Δ CI declined across the distribution (b = −0.050, 95% CI [−0.072, −0.028], t(30) = −4.60, p < .001), confirming that the Simon effect decreased with increasing RT, as it did in the Vincentized raw RT delta functions (Figure S1).

Analogous to mean RT, the COMPATIBILITY × TARGET interaction was restricted to the Δ CV (Δ CI: F(8, 30) = 1.39, p = .242; Δ IV: F(8, 30) = 1.46, p = .212). The Δ CV was larger for tools than for weapons at τ = .10 and .20 (both p ≤ .046) but not at τ = .30 to .90 (all p ≥ .157), and this difference declined from fast to slow responses (b = −0.067, 95% CI [−0.113, −0.022], t(30) = −3.00, p = .005; Table S8). Thus, the cost of vertical relative to compatible trials was larger for tools than for weapons at fast responses but converged at slow responses.

The COMPATIBILITY × CARDIAC PHASE interaction was restricted to the Δ IV (Δ CV: F(8, 30) = 1.60, p = .166; Δ CI: F < 1). The Δ IV was larger during systole than during diastole at τ = .50 and .60 (both p ≤ .015) but not at the remaining quantiles (all p ≥ .053; Table S8), indicating that the cost of vertical relative to incompatible trials was larger during systole at moderate response times. The three-way interaction showed that the phase effect on the Δ CV depended on the prime. Following fear primes, the Δ CV was larger during systole than during diastole at τ = .70 (p = .008) but not at the remaining quantiles (all p ≥ .068), whereas following neutral primes it did not differ between phases at any quantile (F(8, 30) = 1.73, p = .133; all p ≥ .495; Table S8).

Three further interactions reached significance in the omnibus test but could not be localised to any part of the RT distribution, as no per-quantile contrast survived correction across quantiles (COMPATIBILITY × PRIME × TARGET Δ CV: F(8, 30) = 2.28, p = .049, smallest p = .270 at τ = .70; COMPATIBILITY × CONDITION × CARDIAC PHASE × TARGET Δ CI: F(8, 30) = 2.47, p = .035, smallest p = .793 at τ = .80; COMPATIBILITY × CONDITION × CARDIAC PHASE × PRIME × TARGET Δ CI: F(8, 30) = 2.30, p = .047, smallest p = .130 at τ = .40). We therefore do not interpret these interactions further. No further compatibility-involving term reached significance (COMPATIBILITY × CONDITION × CARDIAC PHASE Δ IV: F(8, 30) = 2.18, p = .058; COMPATIBILITY × CARDIAC PHASE × PRIME × TARGET Δ CI: F(8, 30) = 2.16, p = .061; COMPATIBILITY × CONDITION × CARDIAC PHASE × TARGET Δ CV: F(8, 30) = 1.91, p = .096; all remaining terms p ≥ .098).

##### 3.2.3.2. Sensitivity Delta

The CAF GLMM indicated RT-dependent effects on sensitivity of COMPATIBILITY × RT × d′ (χ²(6) = 306.45, p < .001), COMPATIBILITY × RT × CONDITION × d′ (χ²(6) = 25.29, p < .001), and COMPATIBILITY × RT × CARDIAC PHASE × d′ (χ²(6) = 15.35, p = .018).

As expected, RT modulated all three compatibility effects on sensitivity (Δ CI: χ²(3) = 293.12, p < .001; Δ CV: χ²(3) = 57.45, p < .001; Δ IV: χ²(3) = 99.21, p < .001). The Δ CI decreased with increasing RT (b = 1.54, t(7) = 4.37, p = .003, R² = .73), indicating that the Simon effect on sensitivity was largest for fast responses, and the Δ CV followed the same pattern (b = 0.87, t(7) = 11.31, p < .001, R² = .95), whereas the slope of the Δ IV was not significant (b = −0.62, t(7) = −2.33, p = .053, R² = .44).

Decomposing the COMPATIBILITY × RT × CONDITION × d′ interaction showed RT-dependent condition effects on all three contrasts (Δ CI: χ²(3) = 16.75, p < .001; Δ CV: χ²(3) = 13.73, p = .003; Δ IV: χ²(3) = 8.78, p = .032). Relative to control, CPT increased the Δ CI at τ = .10 to .30 (all p ≤ .004) and the Δ CV at τ = .10 (p = .006) and decreased the Δ IV at τ = .20 to .40 (all p ≤ .049), with no condition differences at the remaining quantiles (all p ≥ .127; Table S9A).

Decomposing the COMPATIBILITY × RT × CARDIAC PHASE × d′ interaction showed RT-dependent phase effects on the Δ CV (χ²(3) = 12.24, p = .007) and the Δ CI (χ²(3) = 8.14, p = .043). The Δ CV was larger during systole than during diastole at τ = .10 (p = .026) but not at the remaining quantiles (all p ≥ .061), whereas the Δ CI did not differ between phases at any quantile (all p ≥ .129; Table S9B). No further RT-dependent compatibility effects on sensitivity reached significance (COMPATIBILITY × RT × CONDITION × PRIME × d′: χ²(6) = 11.35, p = .078; all other RT-dependent compatibility χ² < df).

### 3.3. EEG

#### 3.3.1. Midfrontal Theta

Based on the behavioral results, we ran cluster based permutation testing on target-locked midfrontal theta ERSP averaged over 4-8Hz and midfrontal ROI (Fz, Cz, FCz), on the COMPATIBILITY, COMPATIBILITY × TARGET, CONDITION × COMPATIBILITY × TARGET, CONDITION × TARGET × PRIME, CARDIAC PHASE × TARGET × PRIME effects.

As shown in Figure 7 A, the COMPATIBILITY effect on midfrontal theta was qualified by TARGET. For weapon targets the expected Simon effect emerged: theta was higher on incompatible than compatible trials (320 - 896 ms, p = .003, Σt = -535.7) and higher on incompatible than vertical trials (320 - 896 ms, p < .001, Σt = 604.1), whereas compatible and vertical trials did not differ (M = 0.008, 95% CI [-0.117, 0.132], p = 1.000). For tool targets the Simon effect was absent, i.e. no cluster distinguished compatible from incompatible trials (M = -0.015, 95% CI [-0.162, 0.132], p = 1.000), although both compatible and incompatible trials showed higher theta than vertical trials (336 - 624 ms, p = .045, Σt = 186.6; incompatible > vertical: 376 - 680 ms, p = .032, Σt = 215.6). This absent tool Simon effect reflected elevated theta on tool relative to weapon targets: weapon targets elicited less theta than tool targets in compatible (p = .014, Σt = -391.7) and vertical trials (508 - 896 ms, p = .003, Σt = -386.7), whereas weapon and tool targets did not differ in incompatible trials (M = 0.009, 95% CI [-0.115, 0.133], p = 1.000). Formally, however, the target-dependence of the compatibility effect reached significance only for the Δ IV contrast, which differed between targets (M = 0.066, 95% CI [-0.074, 0.206], 560 - 896 ms, p = .026, Σt = 265.3). The compatible-incompatible contrast, i.e. the Simon effect, did not differ significantly between targets (M = -0.165, 95% CI [-0.377, 0.047], p = .064), however since the CIs spans zero yet extends across the full magnitude of the theta difference, we cannot fully exclude a target difference in the theta Simon effect. Finally, the Δ CV contrast likewise did not differ between targets (M = -0.099, 95% CI [-0.256, 0.058], p = 1.000).

The CONDITION × PRIME × TARGET interaction revealed a fear prime specific CONDITION × TARGET effect. Following fear primes, in the CPT condition weapon targets elicited less theta compared to weapon targets under control (392-896 ms, p = .025, Σt = -295.9), and compared to tool targets under CPT (p = .004, Σt = -461.7). Both effects were target- and condition-specific: for tool targets the CPT and control did not differ significantly (M = 0.016, 95% CI [-0.125, 0.158], p = 1.000), and under control weapon and tool clusters were likewise absent (M = 0.017, 95% CI [-0.106, 0.139], p = 1.000). This dissociation was confirmed by a significant CONDITION × TARGET interaction within fear primes, i.e. the weapon–tool difference was more negative under CPT than under control (308 - 700 ms, p = .026, Σt = -253.2). By contrast, following neutral primes, CPT and control did not differ for either weapon (M = -0.021, 95% CI [-0.193, 0.151], p = 1.000) or tool targets (M = -0.027, 95% CI [-0.206, 0.153], p = 1.000), and the CONDITION × TARGET interaction within neutral primes was non-significant (M = 0.006, 95% CI [-0.144, 0.155], p = 1.000).

The PHASE × PRIME × TARGET interaction indicated that cardiac phase shaped the same effect. Following fear primes, during diastole weapon targets elicited less theta than tool in an early (0 - 308 ms, p = .028, Σt = -249.8) and a late window (624 - 896 ms, p = .044, Σt = -198.6), whereas during systole this fear-primed specific weapon-tool contrast was absent (M = 0.002, 95% CI [-0.157, 0.160], p = .176). This dissociation was confirmed by a significant PHASE × TARGET interaction within fear primes over the early window, the weapon-tool contrast being more positive during systole than during diastole (0 - 264 ms, p = .036, Σt = 232.1). Under neutral primes the weapon-tool difference reached significance during systole (452 - 896 ms, p = .022, Σt = -286.7) but not diastole (M = -0.024, 95% CI [-0.153, 0.105], p = 1.000). However, this difference between phases was not supported by an interaction, as neither the PHASE × TARGET interaction within neutral primes (M = -0.073, 95% CI [-0.257, 0.111], p = 1.000) nor the PRIME × TARGET interaction within systole (M = 0.098, 95% CI [-0.087, 0.283], p = 1.000) reached was significant.

#### 3.3.2. Central Beta Lateralization

We ran cluster-based permutation testing on response-locked, response-lateralized LPS in the beta band (13–30 Hz) at a central ROI (FC1, FC3, FC5, C1, C3, C5, CP1, CP3, CP5), on the COMPATIBILITY, COMPATIBILITY × TARGET, CONDITION × COMPATIBILITY × TARGET, CONDITION × PRIME × TARGET, CARDIAC PHASE × PRIME × TARGET, and CARDIAC PHASE × PRIME × COMPATIBILITY effects.

Lateralized beta power differed significantly from zero within every compatibility level. Compatible trials (p < .001, Σt = −2022.8, −484 to 0 ms; channels: CP1, CP3, CP5, C1, C3, C5) and vertical trials (p < .001, Σt = −1403.4, −328 to 0 ms; channels: CP1, CP3, CP5, C1, C3, C5) each yielded a single negative cluster, indicating contralateral suppression building up towards the correct response location. By contrast, incompatible trials showed an early positive cluster (p = .001, Σt = 1210.5, −528 to −192 ms; channels: CP1, CP3, CP5, C1, C3, C5) with significant lateralization towards the incorrect response location succeeded by a negative cluster towards the correct response (p = .002, Σt = −911.1, −132 to 0 ms; channels: CP1, CP3, CP5, C1, C3). Furthermore, compatible trials showed significantly stronger contralateral beta suppression than incompatible trials (M = −0.020, 95% CI [−0.027, −0.013], −552 to −80 ms, p < .001, Σt = −2736.9; channels: CP1, CP3, CP5, C1, C3, C5, FC5, FC3, FC1), and vertical trials showed significantly stronger suppression than incompatible trials (M = 0.015, 95% CI [0.007, 0.022], −524 to −60 ms, p = .002, Σt = 1342.9; channels: CP1, CP3, CP5, C1, C3, FC3, FC1). Compatible trials additionally showed significantly stronger suppression than vertical trials (M = −0.005, 95% CI [−0.012, 0.002]; −348 to −136 ms, p = .032, Σt = −407.5; channels: CP1, CP5, C3, C5, FC5, FC3, FC1). The ordering from strongest to weakest suppression was therefore compatible, vertical, incompatible. This ordering did not significantly deviate between targets, conditions, and none of the corresponding interaction tests reached significance, with interaction estimates centered on zero throughout (COMPATIBILITY × TARGET all p = 1.000, intervals within ±0.013; CONDITION × COMPATIBILITY × TARGET all p ≥ .392; for the full results please refer to Figure S5 in the supplementary materials).

The interaction of CARDIAC PHASE × PRIME × COMPATIBILITY revealed an effect of fear priming confined to systole. On compatible trials, beta suppression was significantly weaker under fear than under neutral primes at systole (M = 0.014, 95% CI [0.002, 0.027], −700 to −88 ms, p = .009, Σt = 689.5; channels: CP3, CP5, C1, C3, C5), but not at diastole (M = 0.001, 95% CI [−0.012, 0.015], p = .525). The direct systole-versus-diastole contrast on compatible trials was not significant under either prime (fear M = 0.004, 95% CI [−0.010, 0.019], p = .540; neutral M = −0.008, 95% CI [−0.023, 0.006], p = .613), and the PHASE × PRIME interaction on compatible trials was likewise not significant (M = 0.013, 95% CI [−0.008, 0.034], p = .519). Because this interval both includes zero and extends beyond the size of the fear effect it would need to explain (0.014), a phase-dependent fear effect can therefore be neither clearly demonstrated nor excluded. No other phase modulation was significant, by either a direct contrast or a formal interaction.

There were no significant clusters in either the cell-wise contrasts or the interactions tests for CONDITION × PRIME × TARGET (all p ≥ .168, all |M| ≤ 0.018, all 95% CIs including zero and no wider than ±0.030) nor PHASE × PRIME × TARGET (all p ≥ .105, all |M| ≤ 0.007, all CIs including zero).

## 4. Discussion

We examined whether cardiac phase and acute stress modulate automatic response tendencies arising from affective and spatial overlap within a common response-selection context. The repeated CPT increased endocrine, cardiovascular, and subjective indices of stress and arousal, replicating earlier work (Larra et al., 2023; Larra et al., 2015; von Haugwitz et al., 2024).

The combined task produced the expected signature of spatial stimulus-response compatibility, i.e. the Simon Effect (Cespon et al., 2020; Hommel, 2011; Simon, 1969). Responses were faster and discrimination sensitivity higher when stimulus and response locations overlapped (e.g., left stimulus – left response) than when they conflicted (e.g., right stimulus – left response, Fig. 3 A; Fig. 4 A), and this effect decreased across the response-time distribution (Fig. 5 A; Fig. 6 A), consistent with a transient influence of automatic spatial response activation (De Jong et al., 1994; Hommel, 1994; Miller & Schwarz, 2021; Ridderinkhof, 2002). Response-locked central beta lateralization showed the corresponding neural signature: compatible trials lateralized toward the correct response, whereas incompatible trials initially lateralized toward the incorrect response before reversing towards the correct response location (Fig. 8 A; c.f. “Gratton dip”; Asanowicz et al., 2021; Gratton et al., 1988). In addition to the two lateral conditions, targets could appear at vertical locations lacking dimensional overlap with the lateralized response set, intended as a neutral reference permitting decomposition of the Simon effect into facilitatory (compatible–vertical, ΔCV) and inhibitory (incompatible–vertical, ΔIV) components (Kornblum et al., 1990; Smith & Ulrich, 2025).

**Fig. 3.**
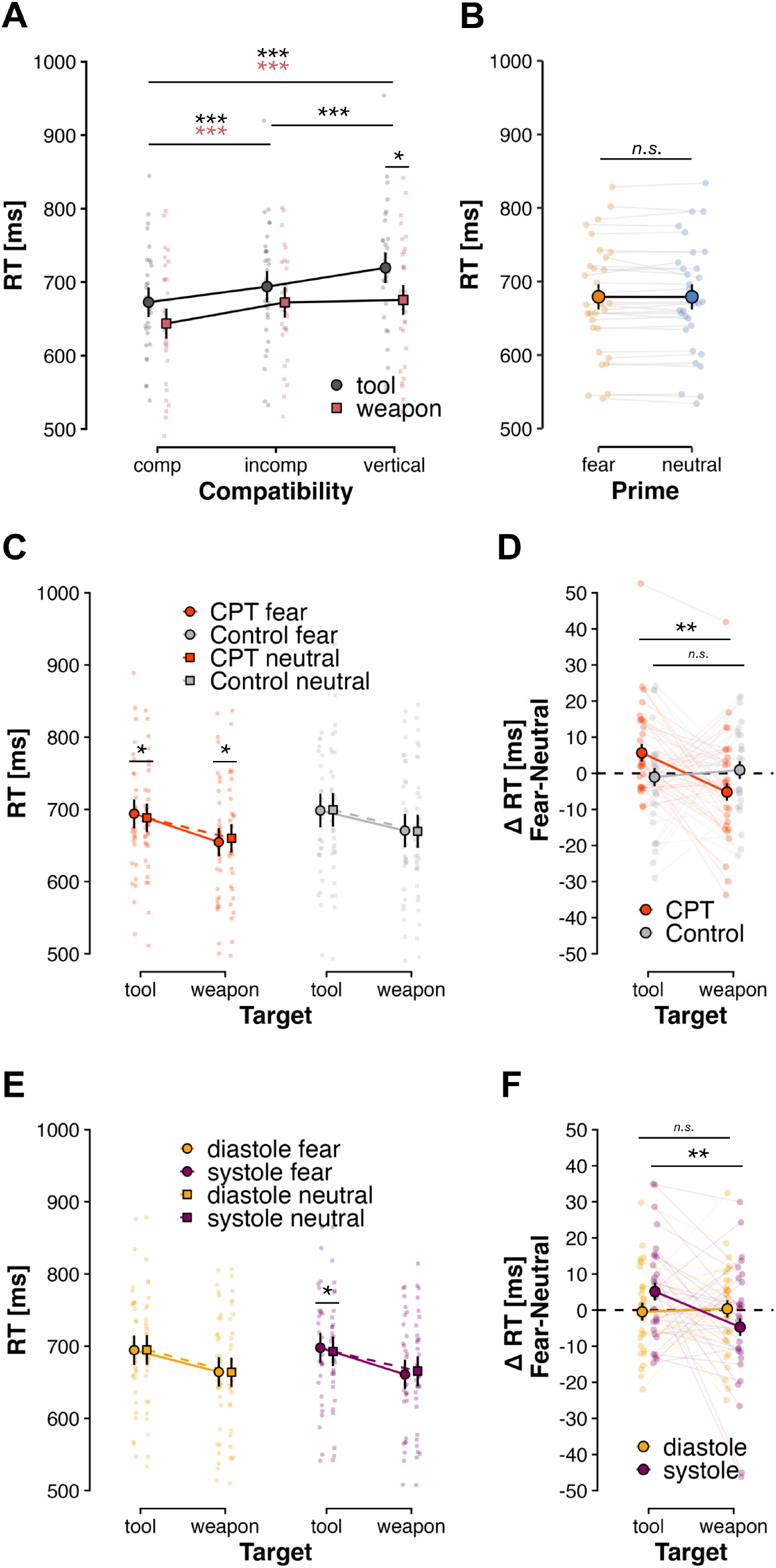
RT Effects of COMPATIBILITY × TARGET (Panel A), PRIME (Panel B), CONDITION × PRIME × TARGET (Panel D and E), and PHASE × PRIME × TARGET (Panel E and F). Error bars show standard errors. *\* p < .05, ** p < .01, *** p < .001*.

**Fig. 4.**
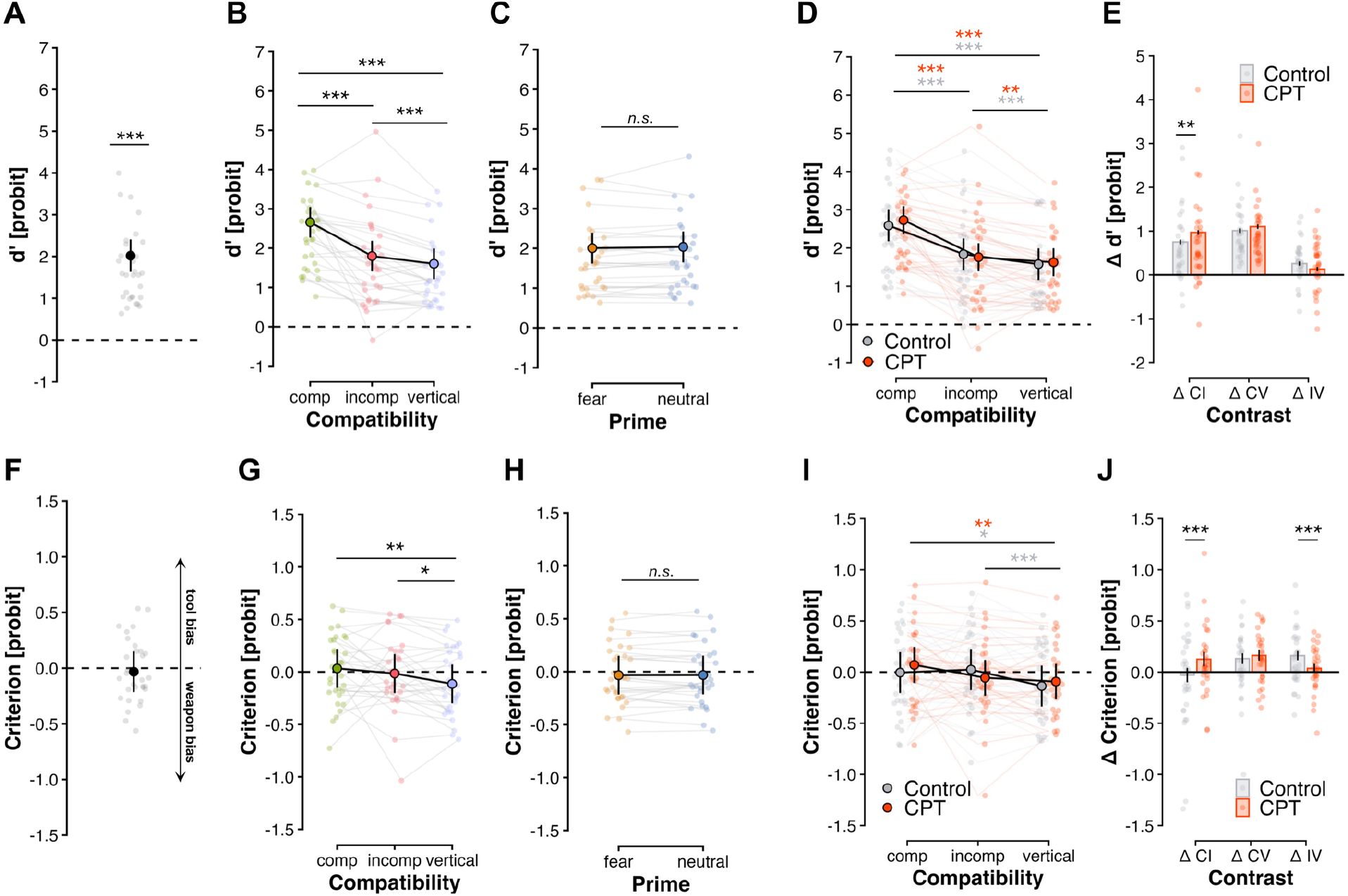
Effects on signal detection theory (SDT) measures. The top row depicts effects on sensitivity (Panels A–E) and the bottom row effects on criterion (Panels F–J). Mean sensitivity and criterion (Panel A and F), effect of COMPATIBILITY (Panel B and G), PRIME (Panel C and H), COMPATIBILITY × CONDITION (sensitivity: Panel D and E; criterion: I and J). Error bars show standard errors. *^n.s.^ p > .05*, *\* p < .05, ** p < .01, *** p < .001*.

**Fig. 5.**
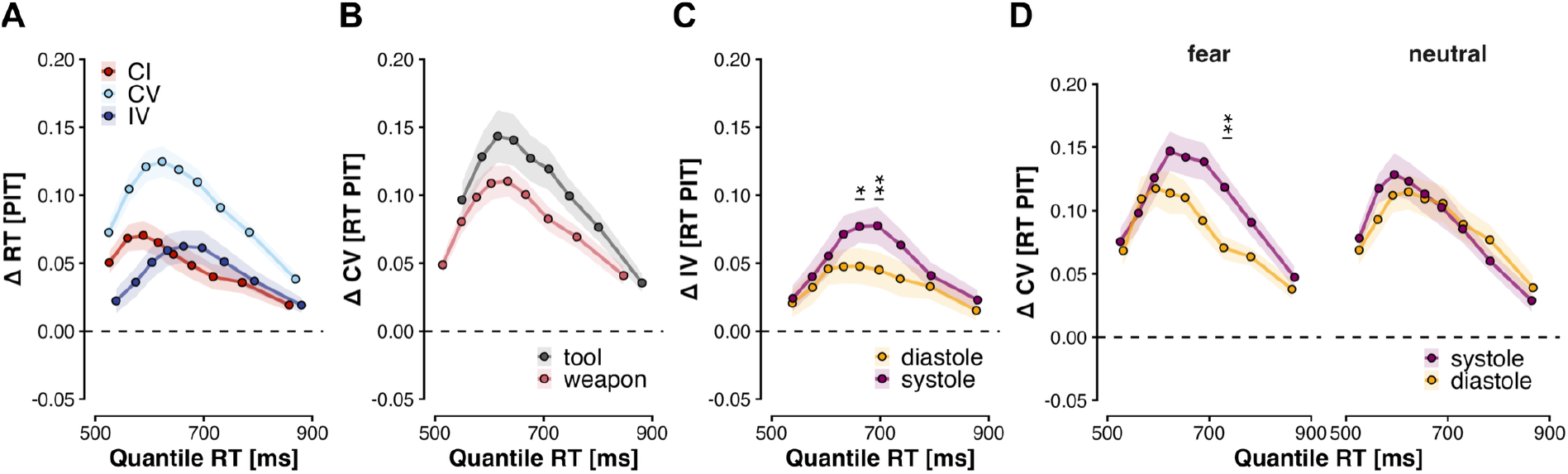
Effect on RT deltas for COMPATIBILITY (Panel A), COMPATIBILITY × TARGET (Panel B), COMPATIBILITY × PHASE (Panel C), COMPATIBILITY × PHASE × PRIME (Panel D). Shaded areas show standard errors. *\* p < .05, ** p < .01*.

**Fig. 6.**
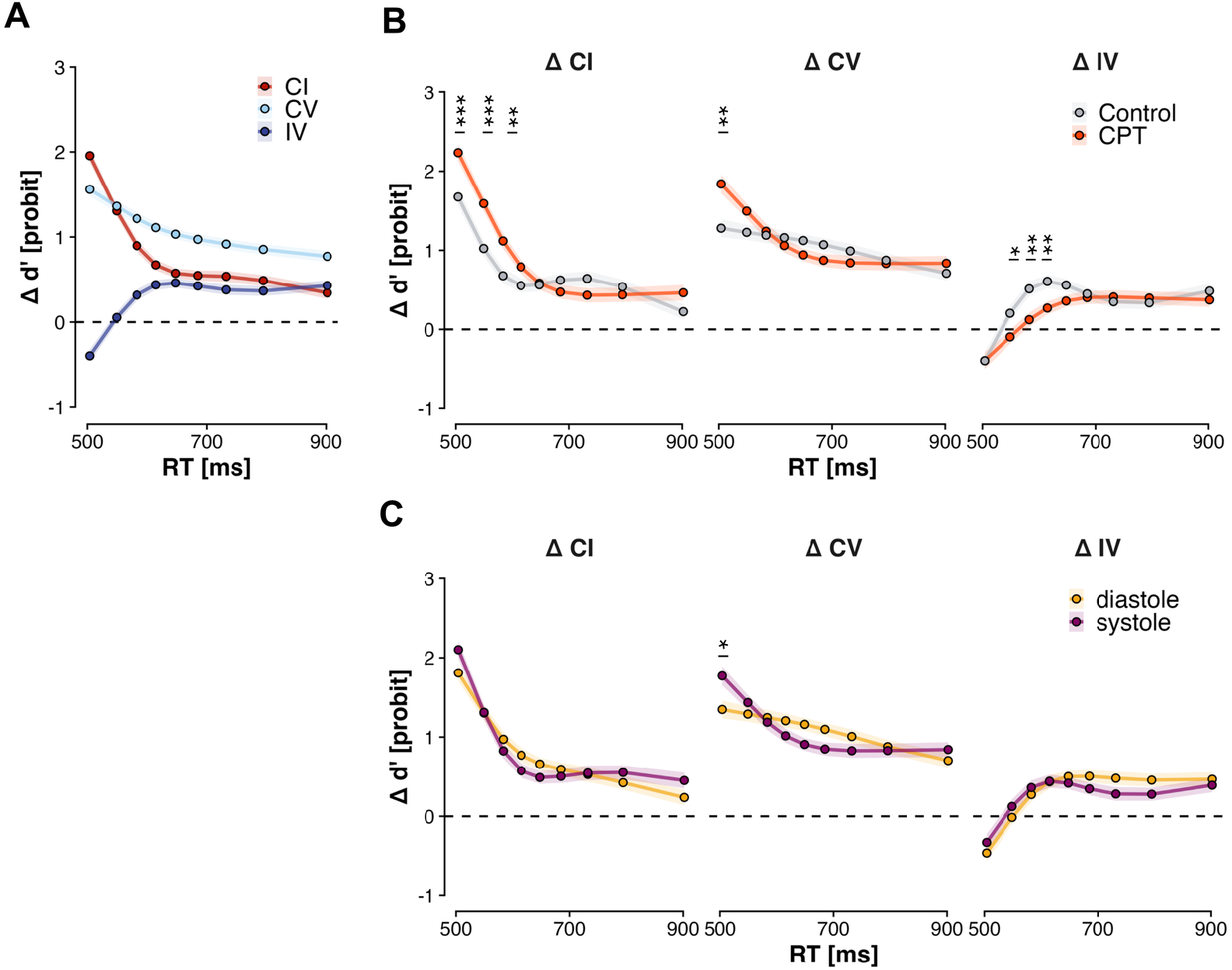
Sensitivity compatibility deltas (Panel A) and effect of CONDITION (Panel B), and PHASE (Panel C) on sensitivity compatibility deltas. Shaded areas show standard errors. *\* p < .05, ** p < .01, *** p < .001*.

**Fig. 7.**
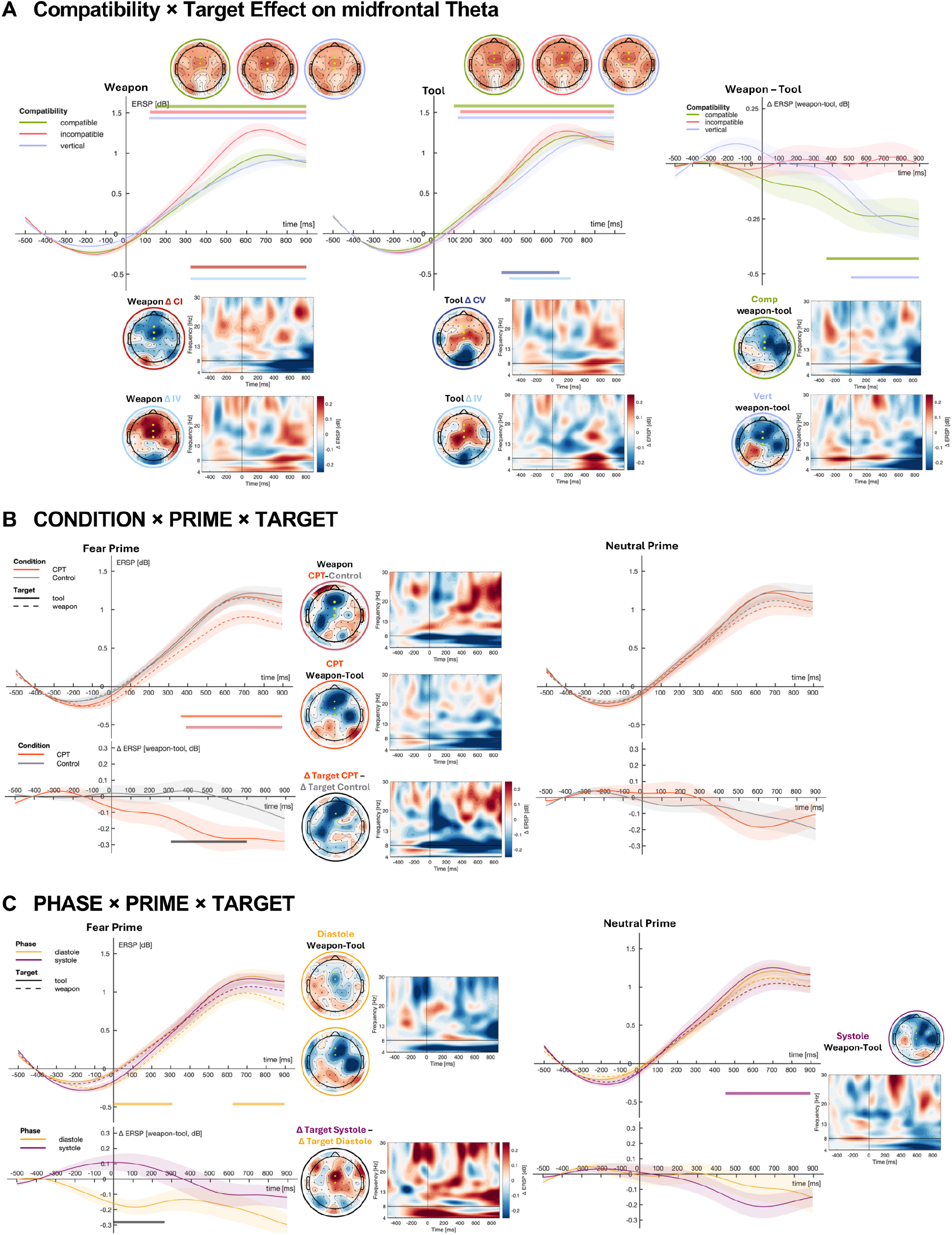
Effect of COMPATIBILTIY × TARGET (Panel A), CONDITION × PRIME × TARGET (Panel B), PHASE × PRIME × TARGET (Panel C) on midfrontal theta (4-8 Hz) at a midfrontal region of interest (Fz, FCz, Cz). Shaded areas around lines represent standard errors. Coloured horizontal bars show significant clusters. Topographies are averaged over significant cluster timepoints. Time-Frequency maps are averaged over the same ROI.

**Fig. 8.**
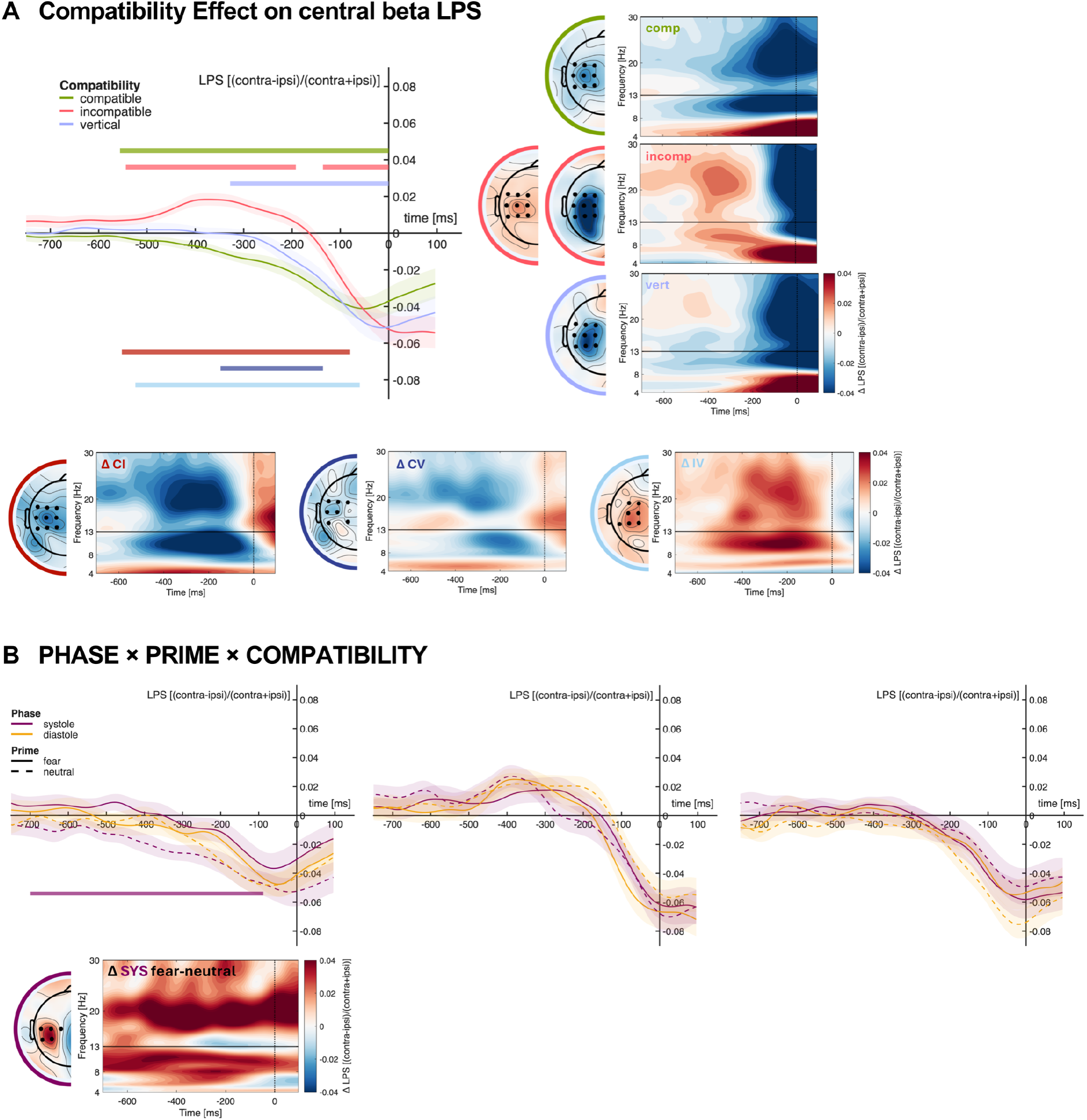
Effect of COMPATIBILTIY (Panel A) and PHASE × PRIME × COMPATIBILITY (Panel B) on beta LPS (13-30 Hz) at a central region of interest (FC1/FC2, FC3/FC4, FC5/FC6, C1/C2, C3/C4, C5/C6, CP1/CP2, CP3/CP4, CP5/CP6). Shaded areas around lines represent standard errors. Coloured horizontal bars show significant clusters. Topographies are averaged over significant cluster timepoints. Time-Frequency maps are averaged over significant cluster channels.

Target category also affected performance most clearly in this vertical non-overlap condition. Weapons were classified faster than tools on vertical trials, particularly among fast responses (Fig. 3 A; Fig. 5 B), accompanied by an increased weapon bias on vertical compared with compatible trials (Fig. 4 G). Target category also modulated the midfrontal theta contrast, with increased theta for tools versus weapons specifically in compatible and vertical trials (Fig. 7 A). This suggests that when the target location primed neither response, intrinsic differences between target categories became more consequential, revealing greater perceptual accessibility of threat-related objects or greater categorization demand for tools (Blanchette, 2006; Brosch & Sharma, 2005; Cavanagh & Frank, 2014; Fox et al., 2007).

Cardiac phase altered how affective context influenced weapon versus tool classification. During systole, prime effects on the two target categories went in opposite directions: tools were classified faster after neutral than fearful primes, whereas weapons showed the complementary advantage after fearful primes while no such interaction emerged during diastole (Fig. 3 E, F). This extends studies using racial primes, in which cardiac phase modulated racially biased weapon identification and shooting decisions (Azevedo et al., 2017; Sherrill et al., 2023) in two ways. First, the effect is not specific to racial stereotypes but also applies to emotional faces, as Azevedo et al. (2017) predicted. Second, we synchronized target onset rather than the prime or a combined cue–target stimulus, showing that cardioafferent signals can bias an already evoked affective context during target categorization or response selection, not only prime encoding. This systolic pattern may reflect threat-directional processing, i.e. fearful primes potentiating weapon processing, with tools acquiring relative safety value (c.f. Azevedo et al., 2017) or a more general strengthening of prime–response correspondence (De Houwer & Eelen, 1998; Hermans et al., 1994). The statistically established simple prime effect concerned tools, consistent with neutral primes producing priming comparable to valenced primes (Scherer & Lambert, 2009) while the weapon advantage was descriptive and did not occur irrespective of prime. The comparable systolic enhancement of Black–weapon and White–tool associations in Azevedo et al. (2017) fits either account, but the absence of modulation for a positive Black–athletic association suggests that systole strengthens prime–response correspondence selectively when the context is affectively salient. In line with this, systolic amplification of emotional faces is not restricted to fear but extends to other expressions, depending on the task (Leganes-Fonteneau et al., 2021).

A further extension is that acute stress also modulated the influence of affective primes. After CPT exposure, fearful compared with neutral primes speeded weapon classification and slowed tool classification while this modulation was absent in the control condition (Fig. 3 C, D). Thus, acute stress increased the influence of task-irrelevant affective context on classification, consistent with a stress-induced shift of processing toward salient, stimulus-driven information (Arnsten, 2009; Hermans et al., 2014). This converges with findings obtained with racial primes, where larger cortisol responses were associated with fewer errors when police officers decided to shoot armed Black relative to armed White targets (Akinola & Mendes, 2012) and with lower process-dissociation estimates of control, that is, a stronger dependence of WIT responses on the prime (Amodio, 2009). The present results extend these correlational findings to an experimentally induced stressor and to emotional primes. As in these studies, stress acted on prime-driven responding rather than on threat responding in general, inducing neither a general bias toward weapon responses nor a deterioration of overall speed or discrimination sensitivity. Rather than prioritizing threat per se, stress appears to have increased the weight of affective context, in line with evidence that acute stress heightens amygdala responses to salient faces at the expense of threat selectivity (van Marle et al., 2009). The theta results further clarify how stress affected control. Under CPT compared with control, theta was reduced for weapons after fearful primes (Fig. 7 B), that is, where behaviorally prime and target converged and responses were faster. An impaired capacity for conflict monitoring, as proposed for stress effects on stereotype control (Amodio, 2009), would instead be expected to manifest primarily on conflicting trials. The reduction therefore more likely reflects a reduced need for control: stress, as the body’s response to threat, may tune processing toward seemingly context-relevant stimuli, so that less control is recruited when the context supports the required response. Together with the preserved overall performance, this suggests a qualitative shift in how control is deployed rather than a uniform loss of executive control (Larra et al., 2022; Plessow et al., 2017)

Cardiac-phase effects on response selection were shaped by spatial context. At moderate RTs, the vertical–incompatible RT difference was larger during systole (Fig. 5C), whereas the compatible–vertical contrast additionally depended on prime, being enhanced during systole after fearful primes (Fig. 5D). In sensitivity, the compatible-vertical difference was likewise larger during systole independent of prime and specific to fast responses (Fig. 6C). A related, prime-dependent effect appeared in response-locked beta lateralization: on compatible trials, fear (relative to neutral) primes were associated with weaker contralateral beta suppression during systole but not diastole (Fig. 8B). Together, these results suggest that cardiac phase acted on the processing advantage conferred by spatial stimulus-response correspondence relative to spatial non-overlap, rather than on response conflict or the classic Simon effect per se, consistent with our earlier work showing that cardiac phase modulates sensorimotor correspondence (Larra et al., 2020; von Haugwitz et al., 2024).

Acute stress also affected spatial response selection. The compatible-incompatible sensitivity difference was larger under CPT than control (Fig. 4 D, E) indicating that stress amplified the spatial compatibility effect, despite neither compatibility condition alone showing a corresponding global stress-related shift. Distributional analyses localized this modulation to fast responses: the compatible-incompatible contrast was enlarged across approximately the fastest third of the response distribution and disappeared at slower responses (Fig. 6 B). Because automatic spatial activation is strongest early and diminishes as responding slows (De Jong et al., 1994; Ridderinkhof, 2002; Ulrich et al., 2015), this pattern points toward stronger initial response activation. Impaired subsequent suppression should instead become increasingly apparent for slower RTs (c.f. Larra et al., 2022; Plessow et al., 2017). This differs from our previous combined horizontal and vertical Simon study, in which acute stress did not modulate behavioral compatibility effects (von Haugwitz et al., 2024). One plausible source of this discrepancy is the conflict context. The previous experiment combined horizontal and vertical Simon effects that are assumed to draw on partly different direct sensorimotor and cognitive routes (Wascher et al., 2001; Wiegand & Wascher, 2005), whereas the present task placed sensorimotor and affective sources of overlap simultaneously onto a single, affectively differentiated response set. Such context may be particularly important under stress. Plessow et al. (2011) found no stress modulation of the overall Simon effect but a reduced congruency-sequence effect, interpreted as increased goal shielding at the expense of flexible context-dependent adjustment. Accordingly, stress may alter how automatic activation is expressed depending on the surrounding conflict structure and on control settings established by other or preceding conflicts, possibly specific to affective ones.

Cardiac phase and acute stress did not interact, for neither the affective nor the spatial effects. This extends previous null findings from affectively neutral tasks (Mussini et al., 2024; von Haugwitz et al., 2024, 2026) to affective processing, where both influences converge, and accords with findings that other state manipulations, such as changes in pupil size or alcohol administration, leave the cardiac modulation of threat-related and emotional processing unchanged (Leganes-Fonteneau, 2024; Sherrill et al., 2023). As outlined in the Introduction, stress could have amplified cardiac effects through its excitatory influence on the amygdala (Garfinkel et al., 2014; Joels & Baram, 2009), or attenuated them through baroreflex suppression (Adlan et al., 2018; Broadley et al., 2005; Gianaros et al., 2012) and an anxiety-like weakening of diastolic inhibition of fear processing (Azevedo et al., 2017; Garfinkel et al., 2014). Such opposing influences may have cancelled each other out, for instance if the vagal rebound that follows stressor offset (Mezzacappa et al., 2001) counteracted stress-related baroreflex suppression. Alternatively, cardiac phase and stress may act on partly distinct components of response selection, as their different theta signatures and spatial effects suggest, and therefore combine additively. However, the absence of a significant interaction does not warrant the conclusion that the two influences are truly independent.

Still, the robustness of these cardiac cycle effects to stress observed here and in previous studies (von Haugwitz et al., 2024, 2026) have relevance beyond the laboratory. Azevedo et al. (2017) motivated cardiac-timing research with split-second decisions in physiologically arousing encounters such as police confrontations, but examined it without sustained arousal. Here, the cardiac modulation of prime effects did not depend on stress, suggesting that it is not abolished in such states. Because it was obtained with emotional rather than racial primes, the facial expressions of people involved in an encounter may, beyond race, bias threat-related decisions, particularly during systole and under stress. In such encounters, heart rate typically increases (Anderson et al., 2002; Baldwin et al., 2019; Bertilsson et al., 2019), which shortens diastole disproportionately relative to systole (Bachani et al., 2025; Wiggers, 1921) and increases the probability that stimuli coincide with systole. Although stimulus onsets outside the laboratory are not time-locked to the cardiac cycle, Lai et al. (2024) recently showed that cardiac-phase effects are not confined to the phase in which a stimulus occurs, as cue timing relative to the cardiac cycle shaped sensorimotor oscillatory activity for more than a second. Thus, even if acute stress leaves the per-stimulus cardiac effect unchanged, it may increase the aggregate influence of cardioafferent signaling on affectively and spatially biased decisions. This influence may be smaller while threat is anticipated but action is not yet possible, as heart rate can then transiently decelerate, a freezing-related bradycardia observed in police recruits anticipating shooting decisions under threat (Hashemi et al., 2019; Hashemi et al., 2021) that turns into tachycardia once action is initiated (Hashemi et al., 2019; Krause et al., 2018). Whether operational experience, which reduces such freezing responses (Ly et al., 2017), shifts this balance is an open question. Because target timing fixed the proportion of systolic and diastolic trials by design, these possibilities could not be addressed here and require studies that assess variations in these proportions as the dependent variable.

Two secondary findings deserve brief consideration. First, acute stress altered the relative criterion differences across spatial conditions. None of the individual criteria differed reliably from zero and no condition showed an absolute stress-induced weapon bias, so this effect should not be interpreted as a general shift toward threat responses (Fig. 4 I, J). Rather, stress altered how spatial context shaped the relative decision criterion: Under control, compatible and incompatible trials showed similar criteria while both were more conservative than vertical trials. Under CPT, the two lateral conditions separated: compatible trials descriptively layed on the conservative and incompatible trials on the liberal side, such that the compatible-incompatible difference was significantly larger than under control, while the incompatible-vertical difference was correspondingly reduced. Thus, among lateral trials, spatial conflict shifted the criterion relatively toward weapon reports under stress, whereas spatial compatibility shifted it toward tool reports. Second, RT varied modestly across the respiratory cycle, with faster responses around inhalation than exhalation, consistent with previous reports of respiratory-phase modulation of sensorimotor processing (Johannknecht & Kayser, 2022; Kluger et al., 2021; Saltafossi et al., 2025).

Some limitations qualify our conclusions. First, vertical trials violated the midpoint assumption of a neutral condition, yielding the slowest RTs and lowest sensitivity rather than falling between compatible and incompatible trials (Fig. 3 A, Fig. 4 A; Smith & Ulrich, 2025). This likely reflects their deliberately reduced probability, making vertical target locations less expected (Druker & Anderson, 2010; Shaw & Shaw, 1977). Electrophysiological data nevertheless suggested that this cost did not arise from response competition: vertical trials showed beta lateralization building toward the correct response, intermediate in magnitude between compatible and incompatible trials and without the initial incorrect lateralization characterizing incompatible trials (Fig. 8 A), while midfrontal theta never exceeded either lateral condition (Fig. 7 A). Thus, although vertical trials preclude a strict quantitative decomposition into Simon facilitation and interference, they retained their function as a response-neutral, spatial non-overlap reference. Second, the mechanistic interpretation of the cardiac-phase theta effect remains indirect. Following fearful primes, weapons elicited less theta than tools during diastole but not during systole (Fig. 7 C), a pattern that fits poorly with a pure conflict-monitoring account. Because midfrontal theta is also sensitive to stimulus salience (Munster et al., 2026), and affective primes can alter the representation of subsequently presented targets (Oh et al., 2025), the cardiac theta pattern may instead reflect a systolic shift toward salience-driven processing (Skora et al., 2022). Third, affective primes did not reliably modulate classification during diastole or under control. The observed behavioral effects can therefore be described as increased influence of the primes under systole and stress, but the design cannot determine whether these manipulations amplified an otherwise weak effect or enabled one to emerge. Moreover, fearful and neutral primes do not cleanly dissociate affective correspondence from threat-directional processing; future studies including primes of opposing valence could separate these accounts more directly.

In conclusion, cardiac phase and acute stress each altered how strongly task-irrelevant affective and spatial information influenced subsequent behavior. Both systolic stimulation and acute stress made affective primes more consequential for weapon/tool classification, without evidence for a general prioritization of threat. Acute stress additionally strengthened the spatial compatible-incompatible sensitivity difference specifically among fast responses, consistent with increased initial automatic response activation, and cardiac-phase effects were present on the advantage conferred by spatial correspondence relative to non-overlapping, vertical locations. Together with their different theta signatures, the findings suggest that phasic cardioafferent signaling and sustained acute stress can converge in increasing the influence of otherwise irrelevant information on action while doing so through partly distinct processes.

## Data and Code Availability

Data and code will be made available upon request after description of research purposes.

## Author Contributions

Leon von Haugwitz: Conceptualization; Data curation; formal analysis; investigation; methodology; visualization; writing – original draft; writing – review and editing. Romina Sadr: Investigation; writing – review and editing. Edmund Wascher: Methodology; supervision; writing – review and editing. Mauro F. Larra: Conceptualization; data curation; funding acquisition; methodology; project administration; supervision; writing – review and editing.

## Use of Generative Artificial Intelligence

The authors used Anthropic’s Claude Opus for two purposes. First, it was used to shorten and proofread the manuscript for syntactic and grammatical errors, particularly to improve the use of English as a second language which was applied to all sections. Second, it was used to debug R and MATLAB code. All suggestions were reviewed and accepted or rejected by the authors. The authors take full responsibility for the content of the manuscript. No sensitive, proprietary, or human-subject data were shared with the tool as only code and non-identifying text were provided.

