## Supplementary Materials for "Stress, Heartbeats, and Biases: Cardiac phase and acute stress modulate affective and spatial response bias"

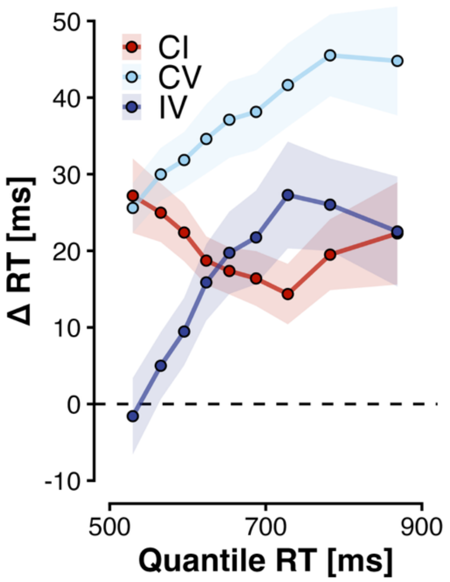


**Figure 1.** Vincentized raw response time delta functions by compatibility delta.

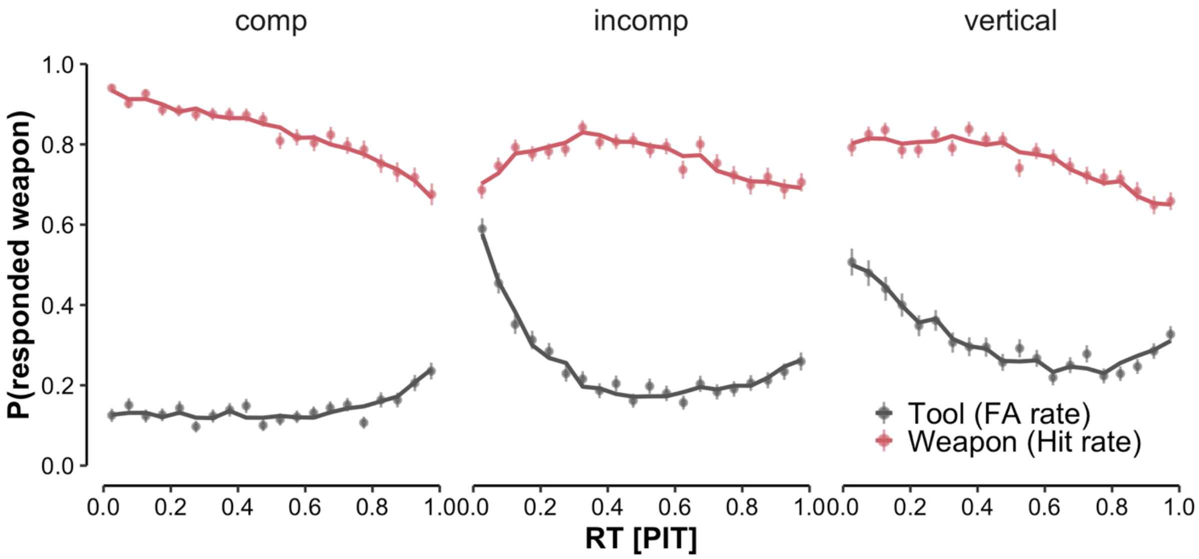


**Figure 2.** Cubic polynomial CAF fit vs. empirical bins. Lines show model predictions while points indicate empirical proportion ± binomial SE.

**Table S1**

*Type III Wald χ² Tests of the Manipulation Check Models.*

| Measure | Effect | $\chi^2$ | *df* | *p* |
|---|---|---|---|---|
| Salivary cortisol | CONDITION | 78.84 | 1 | <.001 |
| | TIME | 63.86 | 6 | <.001 |
| | CONDITION × TIME | 58.47 | 6 | <.001 |
| Mean arterial pressure | CONDITION | 198.93 | 1 | <.001 |
| | TIME | 234.20 | 8 | <.001 |
| | CONDITION × TIME | 229.82 | 8 | <.001 |
| Subjective stress | CONDITION | 128.67 | 1 | <.001 |
| | TIME | 26.32 | 8 | <.001 |
| | CONDITION × TIME | 63.30 | 8 | <.001 |
| Subjective arousal | CONDITION | 211.62 | 1 | <.001 |
| | TIME | 66.07 | 8 | <.001 |
| | CONDITION × TIME | 102.94 | 8 | <.001 |
| Heart rate | CONDITION | 176.57 | 1 | <.001 |
| | TIME | 156.15 | 41 | <.001 |
| | CONDITION × TIME | 209.01 | 41 | <.001 |
| Respiration rate | CONDITION | 44.16 | 1 | <.001 |
| | TIME | 564.19 | 41 | <.001 |
| | CONDITION × TIME | 65.60 | 41 | .009 |

*Note.* Linear mixed models with CONDITION (CPT vs. Control), TIME, and their interaction as fixed effects and a by-subject random intercept.

**Table S2**

Condition x Time Effects on salivary cortisol (nmol/L) throughout the experiment.

| Phase | Bin | Control | | CPT | | Control - CPT | | | |
|---|---|---|---|---|---|---|---|---|---|
| | | *M (SE)* | 95% CI | *M (SE)* | 95% CI | Estimate | *z* | *p* | *d* |
| Rest | 1 | 6.91 (0.87) | [5.22, 8.61] | 5.78 (0.86) | [4.10, 7.47] | 1.13 (0.60) | 1.89 | .069 | 0.18 |
| Block 1 | 2 | 5.82 (0.86) | [4.14, 7.50] | 7.00 (0.86) | [5.32, 8.68] | -1.18 (0.59) | -2.01 | .062 | -0.19 |
| **Block 2** | **3** | **6.03 (0.86)** | **[4.35, 7.71]** | **9.43 (0.86)** | **[7.75, 11.11]** | **-3.40 (0.59)** | **-5.79** | **<.001** | **-0.55** |
| **Block 3** | **4** | **6.22 (0.86)** | **[4.54, 7.90]** | **9.93 (0.86)** | **[8.24, 11.62]** | **-3.71 (0.59)** | **-6.25** | **<.001** | **-0.59** |
| **Block 4** | **5** | **6.08 (0.86)** | **[4.39, 7.77]** | **9.54 (0.86)** | **[7.86, 11.22]** | **-3.46 (0.59)** | **-5.84** | **<.001** | **-0.56** |
| **Post 1** | **6** | **5.38 (0.86)** | **[3.69, 7.06]** | **8.21 (0.86)** | **[6.53, 9.89]** | **-2.83 (0.59)** | **-4.82** | **<.001** | **-0.45** |
| Post 2 | 7 | 5.23 (0.86) | [3.55, 6.91] | 5.65 (0.86) | [3.97, 7.34] | -0.42 (0.59) | -0.72 | .473 | -0.07 |

**Table S3**

*Condition x Time Effects on MAP (mmHg) throughout the experiment.*

| Phase | Bin | Control | | CPT | | Control - CPT | | | |
|---|---|---|---|---|---|---|---|---|---|
| | | *M (SE)* | 95% CI | *M (SE)* | 95% CI | Estimate | *z* | *p* | *d* |
| Rest | 1 | 86.26 (0.99) | [84.32, 88.19] | 85.89 (0.99) | [83.96, 87.83] | 0.37 (0.62) | 0.59 | .554 | 0.06 |
| **CPT 1** | **2** | **86.29 (1.16)** | **[84.01, 88.57]** | **95.58 (1.16)** | **[93.30, 97.86]** | **-9.29 (1.07)** | **-8.68** | **<.001** | **-1.43** |
| Block 1 | 3 | 88.58 (1.16) | [86.30, 90.86] | 87.90 (1.16) | [85.62, 90.19] | 0.68 (1.07) | 0.63 | .554 | 0.10 |
| **CPT 2** | **4** | **87.52 (1.16)** | **[85.23, 89.80]** | **98.81 (1.16)** | **[96.52, 101.09]** | **-11.29 (1.07)** | **-10.54** | **<.001** | **-1.74** |
| Block 2 | 5 | 88.16 (1.16) | [85.88, 90.44] | 89.32 (1.16) | [87.04, 91.60] | -1.16 (1.07) | -1.08 | .417 | -0.18 |
| **CPT 3** | **6** | **86.94 (1.16)** | **[84.65, 89.22]** | **98.00 (1.16)** | **[95.72, 100.28]** | **-11.06 (1.07)** | **-10.33** | **<.001** | **-1.71** |
| Block 3 | 7 | 88.03 (1.16) | [85.75, 90.31] | 89.06 (1.16) | [86.78, 91.35] | -1.03 (1.07) | -0.96 | .431 | -0.16 |
| **CPT 4** | **8** | **86.97 (1.16)** | **[84.69, 89.25]** | **96.42 (1.16)** | **[94.14, 98.70]** | **-9.45 (1.07)** | **-8.83** | **<.001** | **-1.46** |
| Block 4 | 9 | 87.65 (1.16) | [85.36, 89.93] | 89.00 (1.16) | [86.72, 91.28] | -1.35 (1.07) | -1.27 | .370 | -0.21 |

**Table S4**

*Condition x Time Effects on subjective stress (VAS) throughout the experiment.*

| Phase | Bin | Control | | CPT | | Control - CPT | | | |
|---|---|---|---|---|---|---|---|---|---|
| | | *M (SE)* | 95% CI | *M (SE)* | 95% CI | Estimate | *z* | *p* | *d* |
| Rest | 1 | 2.45 (0.48) | [1.50, 3.40] | 2.39 (0.48) | [1.44, 3.34] | 0.06 (0.48) | 0.13 | .893 | 0.02 |
| **CPT 1** | **2** | **1.87 (0.48)** | **[0.92, 2.82]** | **4.97 (0.48)** | **[4.02, 5.92]** | **-3.10 (0.48)** | **-6.46** | **<.001** | **-1.15** |
| Block 1 | 3 | 2.61 (0.48) | [1.66, 3.56] | 3.26 (0.48) | [2.31, 4.21] | -0.65 (0.48) | -1.35 | .201 | -0.24 |
| **CPT 2** | **4** | **1.87 (0.48)** | **[0.92, 2.82]** | **5.26 (0.48)** | **[4.31, 6.21]** | **-3.39 (0.48)** | **-7.06** | **<.001** | **-1.26** |
| Block 2 | 5 | 2.61 (0.48) | [1.66, 3.56] | 3.35 (0.48) | [2.41, 4.30] | -0.74 (0.48) | -1.55 | .157 | -0.28 |
| **CPT 3** | **6** | **2.13 (0.48)** | **[1.18, 3.08]** | **5.29 (0.48)** | **[4.34, 6.24]** | **-3.16 (0.48)** | **-6.59** | **<.001** | **-1.17** |
| **Block 3** | **7** | **2.61 (0.48)** | **[1.66, 3.56]** | **3.74 (0.48)** | **[2.79, 4.69]** | **-1.13 (0.48)** | **-2.35** | **.028** | **-0.42** |
| **CPT 4** | **8** | **2.26 (0.48)** | **[1.31, 3.21]** | **5.35 (0.48)** | **[4.41, 6.30]** | **-3.10 (0.48)** | **-6.46** | **<.001** | **-1.15** |
| **Block 4** | **9** | **2.71 (0.48)** | **[1.76, 3.66]** | **3.84 (0.48)** | **[2.89, 4.79]** | **-1.13 (0.48)** | **-2.35** | **.028** | **-0.42** |

**Table S5**

*Condition x Time Effects on subjective arousal (VAS) throughout the experiment.*

| Phase | Bin | Control | | CPT | | Control - CPT | | | |
|---|---|---|---|---|---|---|---|---|---|
| | | *M (SE)* | 95% CI | *M (SE)* | 95% CI | Estimate | *z* | *p* | *d* |
| Rest | 1 | 2.35 (0.44) | [1.50, 3.21] | 2.65 (0.44) | [1.79, 3.50] | -0.29 (0.46) | -0.63 | .530 | -0.12 |
| **CPT 1** | **2** | **1.97 (0.44)** | **[1.11, 2.83]** | **5.94 (0.44)** | **[5.08, 6.79]** | **-3.97 (0.46)** | **-8.57** | **<.001** | **-1.63** |
| Block 1 | 3 | 2.19 (0.44) | [1.34, 3.05] | 3.10 (0.44) | [2.24, 3.96] | -0.90 (0.46) | -1.95 | .057 | -0.37 |
| **CPT 2** | **4** | **1.84 (0.44)** | **[0.98, 2.70]** | **6.19 (0.44)** | **[5.34, 7.05]** | **-4.35 (0.46)** | **-9.41** | **<.001** | **-1.79** |
| Block 2 | 5 | 2.42 (0.44) | [1.56, 3.28] | 3.35 (0.44) | [2.50, 4.21] | -0.94 (0.46) | -2.02 | .056 | -0.38 |
| **CPT 3** | **6** | **2.03 (0.44)** | **[1.17, 2.89]** | **6.03 (0.44)** | **[5.17, 6.89]** | **-4.00 (0.46)** | **-8.64** | **<.001** | **-1.64** |
| **Block 3** | **7** | **2.42 (0.44)** | **[1.56, 3.28]** | **3.42 (0.44)** | **[2.56, 4.28]** | **-1.00 (0.46)** | **-2.16** | **.046** | **-0.41** |
| **CPT 4** | **8** | **2.35 (0.44)** | **[1.50, 3.21]** | **5.87 (0.44)** | **[5.01, 6.73]** | **-3.52 (0.46)** | **-7.60** | **<.001** | **-1.44** |
| **Block 4** | **9** | **2.48 (0.44)** | **[1.63, 3.34]** | **3.71 (0.44)** | **[2.85, 4.57]** | **-1.23 (0.46)** | **-2.65** | **.015** | **-0.50** |

### Table S6

*Condition x Time Effects on Heart Rate (bpm) throughout the experiment.*

| Phase | Bin | Control | | CPT | | Control - CPT | | | |
|---|---|---|---|---|---|---|---|---|---|
| | | *M* | 95% CI | *M* | 95% CI | Estimate | *z* | *p* | *d* |
| Rest | 1 | 76.08 (1.89) | [72.38, 79.79] | 74.42 (1.90) | [70.70, 78.14] | 1.66 (1.16) | 1.43 | .183 | 0.16 |
| Rest | 2 | 74.62 (1.89) | [70.91, 78.33] | 74.14 (1.90) | [70.42, 77.86] | 0.48 (1.16) | 0.41 | .679 | 0.05 |
| Rest | 3 | 75.94 (1.89) | [72.23, 79.65] | 75.18 (1.90) | [71.46, 78.90] | 0.76 (1.16) | 0.65 | .540 | 0.07 |
| Rest | 4 | 76.43 (1.89) | [72.72, 80.14] | 75.60 (1.90) | [71.88, 79.32] | 0.83 (1.16) | 0.72 | .523 | 0.08 |
| Rest | 5 | 76.06 (1.89) | [72.35, 79.77] | 75.41 (1.90) | [71.69, 79.13] | 0.65 (1.16) | 0.56 | .589 | 0.06 |
| Rest | 6 | 75.66 (1.89) | [71.95, 79.36] | 74.84 (1.90) | [71.13, 78.56] | 0.81 (1.16) | 0.70 | .523 | 0.08 |
| **CPT 1** | **7** | **74.14 (1.89)** | **[70.44, 77.85]** | **78.74 (1.90)** | **[75.03, 82.46]** | **-4.60 (1.16)** | **-3.95** | **<.001** | **-0.44** |
| Block 1 | 8 | 75.57 (1.89) | [71.86, 79.28] | 73.32 (1.90) | [69.60, 77.04] | 2.25 (1.16) | 1.93 | .070 | 0.21 |
| **Block 1** | **9** | **76.64 (1.89)** | **[72.93, 80.35]** | **73.58 (1.90)** | **[69.86, 77.30]** | **3.06 (1.16)** | **2.63** | **.013** | **0.29** |
| **Block 1** | **10** | **77.95 (1.89)** | **[74.24, 81.66]** | **74.00 (1.90)** | **[70.29, 77.72]** | **3.95 (1.16)** | **3.39** | **.002** | **0.37** |
| **Block 1** | **11** | **78.48 (1.89)** | **[74.78, 82.19]** | **75.22 (1.90)** | **[71.50, 78.94]** | **3.26 (1.16)** | **2.80** | **.009** | **0.31** |
| **Block 1** | **12** | **78.90 (1.89)** | **[75.19, 82.60]** | **75.61 (1.90)** | **[71.89, 79.33]** | **3.29 (1.16)** | **2.83** | **.009** | **0.31** |
| **Block 1** | **13** | **79.10 (1.89)** | **[75.39, 82.81]** | **75.45 (1.90)** | **[71.73, 79.16]** | **3.65 (1.16)** | **3.14** | **.004** | **0.35** |
| **Block 1** | **14** | **79.22 (1.89)** | **[75.52, 82.93]** | **75.97 (1.90)** | **[72.25, 79.69]** | **3.26 (1.16)** | **2.80** | **.009** | **0.31** |
| **Block 1** | **15** | **80.56 (1.89)** | **[76.86, 84.27]** | **77.54 (1.90)** | **[73.82, 81.26]** | **3.02 (1.16)** | **2.60** | **.013** | **0.29** |
| **CPT 2** | **16** | **75.18 (1.89)** | **[71.48, 78.89]** | **80.65 (1.90)** | **[76.93, 84.37]** | **-5.47 (1.16)** | **-4.70** | **<.001** | **-0.52** |
| **Block 2** | **17** | **76.77 (1.89)** | **[73.07, 80.48]** | **72.22 (1.90)** | **[68.50, 75.94]** | **4.56 (1.16)** | **3.91** | **<.001** | **0.43** |
| **Block 2** | **18** | **77.83 (1.89)** | **[74.12, 81.53]** | **71.92 (1.90)** | **[68.21, 75.64]** | **5.90 (1.16)** | **5.07** | **<.001** | **0.56** |
| **Block 2** | **19** | **78.44 (1.89)** | **[74.74, 82.15]** | **73.08 (1.90)** | **[69.36, 76.80]** | **5.37 (1.16)** | **4.61** | **<.001** | **0.51** |
| **Block 2** | **20** | **79.06 (1.89)** | **[75.35, 82.77]** | **73.88 (1.90)** | **[70.16, 77.60]** | **5.17 (1.16)** | **4.44** | **<.001** | **0.49** |
| **Block 2** | **21** | **79.34 (1.89)** | **[75.63, 83.05]** | **74.10 (1.90)** | **[70.39, 77.82]** | **5.23 (1.16)** | **4.50** | **<.001** | **0.50** |
| **Block 2** | **22** | **79.26 (1.89)** | **[75.56, 82.97]** | **74.62 (1.90)** | **[70.90, 78.34]** | **4.64 (1.16)** | **3.98** | **<.001** | **0.44** |
| **Block 2** | **23** | **79.62 (1.89)** | **[75.91, 83.32]** | **75.26 (1.90)** | **[71.54, 78.98]** | **4.35 (1.16)** | **3.74** | **<.001** | **0.41** |
| **Block 2** | **24** | **80.15 (1.89)** | **[76.45, 83.86]** | **77.00 (1.90)** | **[73.28, 80.72]** | **3.15 (1.16)** | **2.71** | **.011** | **0.30** |
| **CPT 3** | **25** | **74.11 (1.89)** | **[70.40, 77.82]** | **78.03 (1.90)** | **[74.31, 81.74]** | **-3.92 (1.16)** | **-3.36** | **.002** | **-0.37** |
| **Block 3** | **26** | **76.50 (1.89)** | **[72.80, 80.21]** | **73.24 (1.90)** | **[69.53, 76.96]** | **3.26 (1.16)** | **2.80** | **.009** | **0.31** |
| **Block 3** | **27** | **77.36 (1.89)** | **[73.66, 81.07]** | **73.40 (1.90)** | **[69.68, 77.11]** | **3.97 (1.16)** | **3.41** | **.002** | **0.38** |
| **Block 3** | **28** | **78.60 (1.89)** | **[74.89, 82.31]** | **73.85 (1.90)** | **[70.13, 77.57]** | **4.75 (1.16)** | **4.08** | **<.001** | **0.45** |
| **Block 3** | **29** | **78.84 (1.89)** | **[75.14, 82.55]** | **74.35 (1.90)** | **[70.63, 78.07]** | **4.49 (1.16)** | **3.86** | **<.001** | **0.43** |
| **Block 3** | **30** | **78.23 (1.89)** | **[74.52, 81.94]** | **74.96 (1.90)** | **[71.24, 78.68]** | **3.27 (1.16)** | **2.81** | **.009** | **0.31** |
| **Block 3** | **31** | **78.79 (1.89)** | **[75.08, 82.49]** | **75.66 (1.90)** | **[71.94, 79.38]** | **3.13 (1.16)** | **2.69** | **.011** | **0.30** |
| **Block 3** | **32** | **78.62 (1.89)** | **[74.92, 82.33]** | **75.61 (1.90)** | **[71.89, 79.33]** | **3.02 (1.16)** | **2.59** | **.013** | **0.29** |
| Block 3 | 33 | 78.82 (1.89) | [75.12, 82.53] | 77.05 (1.90) | [73.33, 80.77] | 1.78 (1.16) | 1.52 | .157 | 0.17 |
| **CPT 4** | **34** | **75.39 (1.89)** | **[71.69, 79.10]** | **78.96 (1.90)** | **[75.24, 82.68]** | **-3.56 (1.16)** | **-3.06** | **.005** | **-0.34** |
| **Block 4** | **35** | **75.99 (1.89)** | **[72.28, 79.69]** | **73.17 (1.90)** | **[69.45, 76.89]** | **2.82 (1.16)** | **2.42** | **.021** | **0.27** |
| **Block 4** | **36** | **77.39 (1.89)** | **[73.69, 81.10]** | **73.27 (1.90)** | **[69.55, 76.99]** | **4.12 (1.16)** | **3.54** | **.001** | **0.39** |
| **Block 4** | **37** | **77.25 (1.89)** | **[73.54, 80.96]** | **73.98 (1.90)** | **[70.26, 77.70]** | **3.27 (1.16)** | **2.81** | **.009** | **0.31** |
| **Block 4** | **38** | **77.70 (1.89)** | **[73.99, 81.40]** | **74.09 (1.90)** | **[70.37, 77.81]** | **3.61 (1.16)** | **3.10** | **.005** | **0.34** |
| **Block 4** | **39** | **77.72 (1.89)** | **[74.02, 81.43]** | **74.55 (1.90)** | **[70.84, 78.27]** | **3.17 (1.16)** | **2.72** | **.011** | **0.30** |
| Block 4 | 40 | 77.25 (1.89) | [73.55, 80.96] | 75.85 (1.90) | [72.13, 79.57] | 1.40 (1.16) | 1.20 | .266 | 0.13 |
| Block 4 | 41 | 77.42 (1.89) | [73.71, 81.13] | 75.25 (1.90) | [71.53, 78.97] | 2.17 (1.16) | 1.87 | .079 | 0.21 |
| Block 4 | 42 | 78.11 (1.89) | [74.41, 81.82] | 77.06 (1.90) | [73.34, 80.78] | 1.06 (1.16) | 0.91 | .414 | 0.10 |

**Table S7**

*Condition x Time Effects on Respiration Rate (bpm) throughout the experiment.*

| Phase | Bin | Control | | CPT | | Control - CPT | | | |
|---|---|---|---|---|---|---|---|---|---|
| | | *M* | 95% CI | *M* | 95% CI | Estimate | *z* | *p* | *d* |
| Rest | 1 | 15.51 (0.62) | [14.29, 16.74] | 15.14 (0.63) | [13.91, 16.37] | 0.37 (0.48) | 0.77 | .840 | 0.11 |
| Rest | 2 | 15.48 (0.62) | [14.25, 16.70] | 15.41 (0.63) | [14.18, 16.65] | 0.06 (0.49) | 0.12 | .924 | 0.02 |
| Rest | 3 | 15.91 (0.62) | [14.68, 17.13] | 16.22 (0.63) | [14.99, 17.45] | -0.31 (0.48) | -0.64 | .878 | -0.09 |
| Rest | 4 | 16.29 (0.62) | [15.07, 17.52] | 15.89 (0.63) | [14.66, 17.12] | 0.40 (0.48) | 0.83 | .840 | 0.12 |
| Rest | 5 | 15.89 (0.62) | [14.67, 17.12] | 15.98 (0.63) | [14.75, 17.21] | -0.09 (0.48) | -0.18 | .924 | -0.03 |
| Rest | 6 | 16.47 (0.62) | [15.25, 17.69] | 15.86 (0.63) | [14.63, 17.09] | 0.61 (0.48) | 1.26 | .620 | 0.18 |
| **CPT 1** | **7** | **16.81 (0.62)** | **[15.59, 18.04]** | **14.48 (0.63)** | **[13.25, 15.71]** | **2.33 (0.48)** | **4.82** | **<.001** | **0.67** |
| Block 1 | 8 | 18.27 (0.62) | [17.04, 19.49] | 17.17 (0.63) | [15.94, 18.40] | 1.09 (0.48) | 2.26 | .201 | 0.31 |
| Block 1 | 9 | 18.37 (0.62) | [17.15, 19.60] | 17.47 (0.63) | [16.24, 18.70] | 0.90 (0.48) | 1.87 | .261 | 0.26 |
| Block 1 | 10 | 18.22 (0.62) | [16.99, 19.44] | 17.64 (0.63) | [16.41, 18.87] | 0.58 (0.48) | 1.19 | .655 | 0.17 |
| Block 1 | 11 | 18.30 (0.62) | [17.08, 19.52] | 17.30 (0.63) | [16.07, 18.53] | 1.00 (0.48) | 2.07 | .214 | 0.29 |
| Block 1 | 12 | 18.38 (0.62) | [17.16, 19.60] | 17.41 (0.63) | [16.18, 18.64] | 0.97 (0.48) | 2.00 | .214 | 0.28 |
| Block 1 | 13 | 18.20 (0.62) | [16.97, 19.42] | 17.17 (0.63) | [15.94, 18.39] | 1.03 (0.48) | 2.13 | .214 | 0.30 |
| Block 1 | 14 | 18.08 (0.62) | [16.86, 19.31] | 16.91 (0.63) | [15.68, 18.14] | 1.18 (0.48) | 2.43 | .159 | 0.34 |
| Block 1 | 15 | 17.52 (0.62) | [16.29, 18.74] | 16.84 (0.63) | [15.61, 18.07] | 0.68 (0.48) | 1.41 | .607 | 0.20 |
| **CPT 2** | **16** | **15.92 (0.62)** | **[14.70, 17.15]** | **13.55 (0.63)** | **[12.32, 14.78]** | **2.38 (0.48)** | **4.91** | **<.001** | **0.68** |
| Block 2 | 17 | 17.52 (0.62) | [16.30, 18.75] | 17.26 (0.63) | [16.03, 18.49] | 0.26 (0.48) | 0.54 | .916 | 0.08 |
| Block 2 | 18 | 17.84 (0.62) | [16.61, 19.06] | 17.21 (0.63) | [15.98, 18.44] | 0.63 (0.48) | 1.30 | .620 | 0.18 |
| Block 2 | 19 | 17.84 (0.62) | [16.61, 19.06] | 17.73 (0.63) | [16.50, 18.96] | 0.11 (0.48) | 0.22 | .924 | 0.03 |
| Block 2 | 20 | 17.47 (0.62) | [16.25, 18.70] | 17.38 (0.63) | [16.15, 18.61] | 0.09 (0.48) | 0.19 | .924 | 0.03 |
| Block 2 | 21 | 17.47 (0.62) | [16.25, 18.69] | 17.27 (0.63) | [16.04, 18.50] | 0.20 (0.48) | 0.41 | .924 | 0.06 |
| Block 2 | 22 | 17.66 (0.62) | [16.43, 18.88] | 17.24 (0.63) | [16.01, 18.47] | 0.42 (0.48) | 0.86 | .840 | 0.12 |
| Block 2 | 23 | 17.75 (0.62) | [16.53, 18.98] | 17.13 (0.63) | [15.90, 18.36] | 0.62 (0.48) | 1.28 | .620 | 0.18 |
| Block 2 | 24 | 17.17 (0.62) | [15.95, 18.39] | 16.83 (0.63) | [15.60, 18.06] | 0.34 (0.48) | 0.69 | .854 | 0.10 |
| **CPT 3** | **25** | **15.81 (0.62)** | **[14.59, 17.04]** | **13.95 (0.63)** | **[12.72, 15.18]** | **1.86 (0.48)** | **3.84** | **.002** | **0.54** |
| Block 3 | 26 | 17.35 (0.62) | [16.12, 18.57] | 16.81 (0.63) | [15.58, 18.05] | 0.54 (0.49) | 1.10 | .688 | 0.15 |
| Block 3 | 27 | 17.62 (0.62) | [16.40, 18.85] | 17.24 (0.63) | [16.01, 18.47] | 0.39 (0.48) | 0.80 | .840 | 0.11 |
| Block 3 | 28 | 17.64 (0.62) | [16.42, 18.87] | 17.31 (0.63) | [16.08, 18.53] | 0.34 (0.48) | 0.70 | .854 | 0.10 |
| Block 3 | 29 | 17.39 (0.62) | [16.17, 18.62] | 17.30 (0.63) | [16.07, 18.53] | 0.10 (0.48) | 0.20 | .924 | 0.03 |
| Block 3 | 30 | 17.22 (0.62) | [16.00, 18.45] | 17.30 (0.63) | [16.07, 18.53] | -0.08 (0.48) | -0.16 | .924 | -0.02 |
| Block 3 | 31 | 17.16 (0.62) | [15.94, 18.39] | 17.26 (0.63) | [16.03, 18.49] | -0.10 (0.48) | -0.20 | .924 | -0.03 |
| Block 3 | 32 | 17.36 (0.62) | [16.13, 18.58] | 17.53 (0.63) | [16.30, 18.76] | -0.17 (0.48) | -0.36 | .924 | -0.05 |
| Block 3 | 33 | 16.84 (0.62) | [15.62, 18.07] | 16.55 (0.63) | [15.32, 17.78] | 0.29 (0.48) | 0.60 | .886 | 0.08 |
| CPT 4 | 34 | 15.42 (0.62) | [14.19, 16.64] | 14.43 (0.63) | [13.20, 15.66] | 0.99 (0.48) | 2.05 | .214 | 0.28 |
| Block 4 | 35 | 17.28 (0.62) | [16.06, 18.51] | 17.21 (0.63) | [15.99, 18.44] | 0.07 (0.48) | 0.14 | .924 | 0.02 |
| Block 4 | 36 | 17.59 (0.62) | [16.36, 18.81] | 17.19 (0.63) | [15.96, 18.41] | 0.40 (0.48) | 0.83 | .840 | 0.12 |
| Block 4 | 37 | 17.71 (0.62) | [16.49, 18.94] | 17.19 (0.63) | [15.96, 18.42] | 0.52 (0.48) | 1.08 | .688 | 0.15 |
| Block 4 | 38 | 17.75 (0.62) | [16.53, 18.97] | 17.57 (0.63) | [16.34, 18.80] | 0.18 (0.48) | 0.38 | .924 | 0.05 |
| Block 4 | 39 | 17.34 (0.62) | [16.11, 18.56] | 17.49 (0.63) | [16.26, 18.72] | -0.15 (0.48) | -0.32 | .924 | -0.04 |
| Block 4 | 40 | 17.48 (0.62) | [16.26, 18.71] | 17.35 (0.63) | [16.12, 18.58] | 0.14 (0.48) | 0.28 | .924 | 0.04 |
| Block 4 | 41 | 17.31 (0.62) | [16.09, 18.53] | 17.31 (0.63) | [16.08, 18.54] | -0.00 (0.48) | -0.00 | .998 | -0.00 |
| Block 4 | 42 | 16.47 (0.62) | [15.24, 17.69] | 16.59 (0.63) | [15.35, 17.83] | -0.12 (0.49) | -0.25 | .924 | -0.04 |

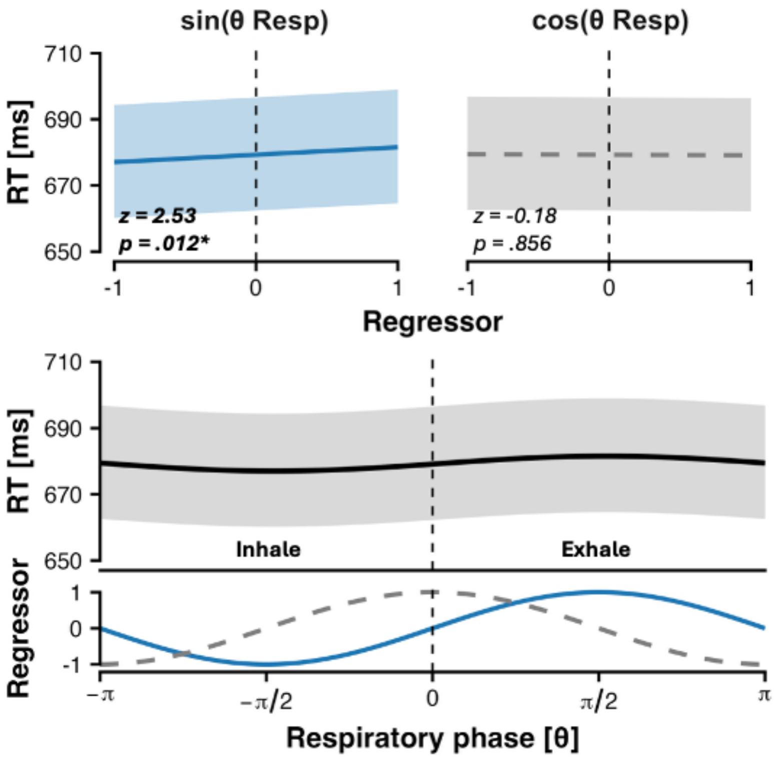


**Figure 3.** Effect of sine and cosine components of respiratory phase on reaction time (RT).

**Table S8**

*Target and Cardiac Phase Effects on the RT Compatibility Deltas per Quantile*

| | *COMP x TARGET* | | | *COMP x PHASE* | | |
|---|---|---|---|---|---|---|
| τ | **Δ CV, tool – weapon** | ***t*** | ***p*** | **Δ IV, sys – dia** | ***t*** | ***p*** |
| .10 | **−0.048 (0.014)** | **−3.52** | **.013** | −0.004 (0.008) | −0.45 | .657 |
| .20 | **−0.048 (0.017)** | **−2.74** | **.046** | −0.008 (0.010) | −0.79 | .488 |
| .30 | −0.045 (0.022) | −2.02 | .157 | −0.009 (0.010) | −0.95 | .488 |
| .40 | −0.032 (0.023) | −1.38 | .402 | −0.024 (0.010) | −2.30 | .065 |
| .50 | −0.017 (0.021) | −0.79 | .559 | **−0.029 (0.009)** | **−3.18** | **.015** |
| .60 | −0.019 (0.018) | −1.03 | .480 | **−0.033 (0.009)** | **−3.67** | **.008** |
| .70 | −0.017 (0.017) | −1.01 | .480 | −0.025 (0.010) | −2.51 | .053 |
| .80 | −0.007 (0.014) | −0.51 | .615 | −0.008 (0.010) | −0.81 | .488 |
| .90 | 0.005 (0.009) | 0.62 | .611 | −0.008 (0.009) | −0.84 | .488 |
| | *COMPATIBILITY × CARDIAC PHASE × PRIME (Δ CV, systole − diastole)* | | | | | |
| | **Fear** | ***t*** | ***p*** | **Neutral** | ***t*** | ***p*** |
| .10 | −0.007 (0.013) | −0.54 | .653 | −0.009 (0.011) | −0.89 | .796 |
| .20 | 0.011 (0.018) | 0.61 | .653 | −0.025 (0.012) | −2.00 | .495 |
| .30 | −0.008 (0.018) | −0.45 | .653 | −0.017 (0.017) | −0.99 | .796 |
| .40 | −0.033 (0.021) | −1.62 | .210 | −0.008 (0.016) | −0.52 | .871 |
| .50 | −0.032 (0.018) | −1.82 | .178 | −0.004 (0.017) | −0.21 | .871 |
| .60 | −0.047 (0.019) | −2.51 | .068 | 0.003 (0.017) | 0.16 | .871 |
| .70 | **−0.048 (0.013)** | **−3.68** | **.008** | 0.004 (0.016) | 0.24 | .871 |
| .80 | −0.027 (0.011) | −2.40 | .068 | 0.017 (0.014) | 1.18 | .796 |
| .90 | −0.009 (0.011) | −0.89 | .571 | 0.010 (0.013) | 0.78 | .796 |

*Note.* Estimates (SE) are differences in the compatibility deltas of PIT-transformed RT at each quantile τ, with Δ CV = compatible – vertical and Δ IV = incompatible − vertical. Negative values therefore indicate a larger cost of vertical trials for the first-named level. *p* values are Benjamini–Hochberg corrected across quantiles.

**Table S9**

*Condition and Cardiac Phase Effects on the Sensitivity Compatibility Deltas per RT Quantile.*

| | Δ CI | | | Δ CV | | | Δ IV | | |
|---|---|---|---|---|---|---|---|---|---|
| τ | *M* | *z* | *p* | *M* | *z* | *p* | M | *z* | *p* |
| | | | | *A: CPT vs. Control* | | | | | |
| .10 | **0.56 (0.15)** | **3.68** | **.001** | **0.56 (0.16)** | **3.42** | **.006** | 0.00 (0.15) | 0.00 | .996 |
| .20 | **0.57 (0.13)** | **4.32** | **<.001** | 0.27 (0.13) | 2.00 | .204 | **−0.30 (0.12)** | **−2.41** | **.049** |
| .30 | **0.44 (0.14)** | **3.24** | **.004** | 0.05 (0.14) | 0.36 | .734 | **−0.39 (0.13)** | **−3.06** | **.018** |
| .40 | 0.24 (0.12) | 1.91 | .127 | −0.10 (0.13) | −0.79 | .550 | **−0.34 (0.12)** | **−2.88** | **.018** |
| .50 | 0.02 (0.11) | 0.17 | .862 | −0.18 (0.11) | −1.62 | .238 | −0.20 (0.11) | −1.90 | .129 |
| .60 | −0.15 (0.12) | −1.26 | .269 | −0.20 (0.12) | −1.70 | .238 | −0.05 (0.11) | −0.43 | .754 |
| .70 | −0.21 (0.13) | −1.62 | .157 | −0.15 (0.13) | −1.19 | .420 | 0.06 (0.12) | 0.51 | .754 |
| .80 | −0.10 (0.13) | −0.81 | .469 | −0.04 (0.12) | −0.34 | .734 | 0.06 (0.12) | 0.52 | .754 |
| .90 | 0.24 (0.14) | 1.72 | .153 | 0.13 (0.13) | 0.97 | .500 | −0.11 (0.13) | −0.89 | .669 |
| | | | | *B: Systole vs. Diastole* | | | | | |
| .10 | 0.29 (0.15) | 1.93 | .129 | **0.43 (0.16)** | **2.60** | **.026** | | | |
| .20 | 0.01 (0.13) | 0.06 | .998 | 0.15 (0.13) | 1.09 | .520 | | | |
| .30 | −0.15 (0.14) | −1.08 | .524 | −0.06 (0.14) | −0.42 | .906 | | | |
| .40 | −0.20 (0.12) | −1.59 | .248 | −0.19 (0.13) | −1.52 | .282 | | | |
| .50 | −0.17 (0.11) | −1.50 | .289 | −0.25 (0.11) | −2.26 | .061 | | | |
| .60 | −0.09 (0.12) | −0.74 | .738 | −0.25 (0.12) | −2.15 | .081 | | | |
| .70 | 0.02 (0.13) | 0.15 | .987 | −0.18 (0.13) | −1.43 | .326 | | | |
| .80 | 0.13 (0.12) | 1.04 | .552 | −0.05 (0.12) | −0.41 | .913 | | | |
| .90 | 0.22 (0.14) | 1.55 | .266 | 0.14 (0.13) | 1.07 | .534 | | | |

*Note*. Estimates (SE) are differences in the *d*' compatibility deltas at each RT quantile. *p* values are multivariate-t adjusted across quantiles.

### Stimulus-locked frontocentral theta: CONDITION x COMPATIBILITY x TARGET

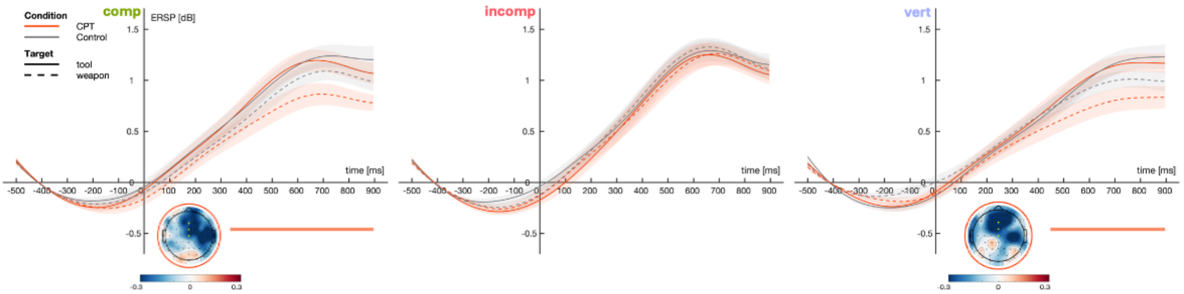


**Figure 4.** COND x COMP x TARGET effect.

The CONDITION × COMPATIBILITY × TARGET analysis showed CPT-specific simple effects that were not, however, confirmed by any direct between-condition test. In both conditions the Simon effect on midfrontal theta was confined to weapon targets, with incompatible trials exceeding both compatible (CPT: M = -0.185, 95% CI [-0.333, -0.036], 364–896 ms, p = .005, Σt = -521.8; Control: M = -0.175, 95% CI [-0.345, -0.004], 356–776 ms, p = .023, Σt = -263.8) and vertical trials (CPT: M = 0.211, 95% CI [0.047, 0.376], 352–896 ms, p = .002, Σt = 530.4; Control: M = 0.163, 95% CI [0.029, 0.298], 360–800 ms, p = .003, Σt = 423.3). Under CPT, the reduction of theta to weapon relative to tool targets extended beyond incompatible trials (where it was absent, M = -0.002, 95% CI [-0.163, 0.158], p = 1.000) to compatible (M = -0.203, 95% CI [-0.339, -0.068], 336–896 ms, p = .001, Σt = -534.1) and vertical trials (M = -0.135, 95% CI [-0.297, 0.027], 448–896 ms, p = .003, Σt = -433.8), and within CPT the target effect varied across compatibility levels (compatible vs incompatible: M = -0.201, 95% CI [-0.425, 0.024], 480–896 ms, p = .027, Σt = -259.6; incompatible vs vertical: M = 0.133, 95% CI [-0.084, 0.349], 548–896 ms, p = .019, Σt = 284.5; compatible vs vertical: M = -0.068, 95% CI [-0.275, 0.139], p = 1.000). Under control, by contrast, the weapon−tool difference was weaker and non-significant on compatible trials (M = -0.110, 95% CI [-0.301, 0.082], p = 1.000) and absent on vertical trials (M = 0.020, 95% CI [-0.116, 0.156], p = .104), and none of the within-control target-by-compatibility interactions reached significance (all p ≥ .177). Crucially, however, the direct between-condition comparisons provided no evidence that stress altered midfrontal theta at any cell: CPT did not differ from control for weapon targets on compatible (M = -0.086, 95% CI [-0.276, 0.104], p = .091), incompatible (M = -0.076, 95% CI [-0.271, 0.118], p = 1.000), or vertical trials (M = -0.124, 95% CI [-0.301, 0.052], p = 1.000), nor for tool targets on any compatibility level (compatible: M = 0.008, 95% CI [-0.160, 0.176], p = 1.000; incompatible: M = -0.054, 95% CI [-0.241, 0.133], p = 1.000; vertical: M = 0.030, 95% CI [-0.159, 0.220], p = 1.000). Consistent with this, the weapon−tool delta did not differ between conditions on compatible (M = -0.094, 95% CI [-0.283, 0.095], p = 1.000), incompatible (M = -0.022, 95% CI [-0.232, 0.187], p = 1.000), or vertical trials (M = -0.155, 95% CI [-0.362, 0.052], p = 1.000), and the three-way tests of whether the COMPATIBILITY × TARGET interaction itself differed between conditions were likewise non-significant for every compatibility pair (incompatible vs vertical: M = 0.133, 95% CI [-0.164, 0.429], p = .141; compatible vs incompatible: M = -0.071, 95% CI [-0.382, 0.239], p = 1.000; compatible vs vertical: M = 0.061, 95% CI [-0.259, 0.381], p = 1.000). The CPT-specific pattern therefore rests on within-condition differences in significance rather than on any demonstrated between-condition difference, and we do not interpret it as a genuine modulation of the compatibility–target relationship by stress.

#### Response-locked central beta LPS: COMPATIBILITY x TARGET; CONDITION x COMPATIBILITY x TARGET

None of the three Compatibility × Target interaction tests approached significance (compatible–incompatible $p = 1.000$, $M = 0.002$, 95% CI [−0.011, 0.014]; compatible–vertical $p = 1.000$, $M = 0.004$, 95% CI [−0.008, 0.015]; incompatible–vertical $p = 1.000$, $M = 0.002$, 95% CI [−0.010, 0.014]), despite the cell-wise contrasts reaching significance in different cells. The two contrasts involving incompatible trials held for both target categories: compatible trials showed significantly stronger suppression than incompatible for weapon ($p < .001$, $\Sigma t = -1455.8$, −512 to −64 ms; channels: CP1, CP3, CP5, C1, C3, C5, FC5, FC1; $M = -0.019$, 95% CI [−0.028, −0.009]) and for tool ($p = .001$, $\Sigma t = -1808.5$, −580 to −108 ms; channels: CP1, CP3, CP5, C1, C3, C5, FC5, FC3, FC1; $M = -0.021$, 95% CI [−0.032, −0.009]), and vertical trials showed significantly stronger suppression than incompatible for weapon ($p < .001$, $\Sigma t = 1015.5$, −532 to −56 ms; channels: CP1, CP3, C1, C3, C5, FC1; $M = 0.016$, 95% CI [0.006, 0.025]) and for tool ($p = .018$, $\Sigma t = 654.2$, −412 to −76 ms; channels: CP1, CP3, CP5, C1, C3, FC3, FC1; $M = 0.014$, 95% CI [0.002, 0.026]). The compatible-versus-vertical contrast, significant across targets, did not reach significance within either target separately (weapon $p = .898$; tool $p = .102$). Weapon and tool did not differ in absolute LPS at any compatibility level (all $p \geq .460$). The compatibility ordering is therefore best described as target-invariant. CONDITION × COMPATIBILITY × TARGET reproduced the same ordering within both conditions, with the individual contrasts reaching significance in different cells, but none of the family's nineteen interaction tests was significant (all $p \geq .392$; full statistics in Supplementary Table S1). Compatible trials showed significantly stronger suppression than incompatible under Control for weapon ($p = .001$, $\Sigma t = -1069.3$, −492 to −16 ms; channels: CP1, CP3, CP5, C1, C3, C5, FC5, FC3, FC1; $M = -0.024$, 95% CI [−0.036, −0.012]) and tool ($p = .017$, $\Sigma t = -675.2$, −384 to −140 ms; channels: CP1, CP3, C1, C3, FC5, FC3, FC1; $M = -0.017$, 95% CI [−0.032, −0.001]), and under CPT for tool ($p = .002$, $\Sigma t = -1329.8$, −504 to −88 ms; channels: CP1, CP3, CP5, C1, C3, C5, FC5, FC3, FC1; M =

−0.025, 95% CI [−0.039, −0.011]); the same contrast did not reach significance under CPT for weapon ($p$ = .082). Vertical trials showed significantly stronger suppression than incompatible under CPT for both tool ($p$ = .023, Σt = 514.3, −436 to −8 ms; channels: CP1, CP3, C1, FC1) and weapon ($p$ = .023, Σt = 536.8, −412 to −112 ms; channels: CP1, CP3, C1, C3, C5, FC1; M = 0.015, 95% CI [0.001, 0.030]), and under Control for weapon ($p$ = .038, Σt = 381.8, −460 to −148 ms; channels: CP1, CP3, C1, C3, C5; M = 0.016, 95% CI [0.003, 0.029]) but not tool ($p$ = .324). Compatible versus vertical did not reach significance in any of the four condition-by-target cells (all $p \geq .191$). No direct CPT-versus-Control comparison was significant at any fixed compatibility-by-target combination (all $p \geq .364$), and weapon and tool did not differ in any cell (all $p \geq .423$). The beta data therefore provide no evidence that acute stress altered the lateralization signature of spatial compatibility.
